# The trimeric structures of the extracellular domains of FAM171A1 and FAM171A2 neuronal proteins belong to a novel structural superfamily

**DOI:** 10.1101/2025.09.18.675241

**Authors:** Tom Bird, Sepideh Valimehr, David M. Wood, Zachary D. Tillett, Leanid Kresik, Gerd Mittelstädt, Florian De Pol, Dimphna H. Meijer, Renwick C.J. Dobson, Joris de Wit, Eric Hanssen, Davide Comoletti

## Abstract

Cell surface molecules play fundamental roles in cell-cell communication, attraction, or repulsion, and when expressed in neurons they are often implicated in neurological disorders. FAM171 is a family of three type-I transmembrane domain cell surface proteins (FAM171A1, FAM171A2, and FAM171B) expressed in several human tissues and especially enriched in the brain. Recent findings suggest that FAM171A1 transduces signals between the cell surface and the cytoskeleton. Genetic evidence links FAM171A1 to multiple cancers and FAM171A2 to neurodegenerative diseases, including Alzheimer’s and Parkinson’s diseases. Despite multiple connections with severe human diseases, no information is currently available on their monomeric structure or oligomerization. Here we show that, structurally, the monomeric ectodomains of human FAM171A1 and FAM171A2 have a new architecture with a novel combination of two domains. Furthermore, their ectodomains oligomerize to form an equilateral trimer. In addition, the ectodomain of FAM171A1 has the propensity to form larger trimer-trimer assemblies at high concentrations. Together, these results provide novel insights into the structure and oligomerization of the extracellular domain of FAM171A1 and FAM171A2, suggesting important roles in ligand binding and signaling.

## INTRODUCTION

FAM171 is an understudied family of cell surface type-I receptors consisting of three paralogs - FAM171A1, FAM171A2 and FAM171B. FAM171A1 is abundantly expressed in the heart, brain, liver, skeletal muscle, kidney, and pancreas^1^. In the brain, FAM171A1 expression is particularly robust in astrocytes (hence the alternative name Astroprincin) and pyramidal neurons. Early investigations of FAM171A1 function describe it to have a regulatory role in cell shape and motility^1^. Overexpression of FAM171A1 in several cell lines induces sprouting of neurite-like projections, suggesting that FAM171A1 transduces signals from the cell surface to the cytoskeleton. FAM171A1 is also able to induce invasive tumor growth, whereas truncation of the C-terminal 21 amino acids ablates this activity^1^. Furthermore, FAM171A1 has been identified as a potential marker and therapeutic target for multiple cancers, and in triple negative breast cancer (TNBC), higher FAM171A1 expression levels are associated with poor prognosis^2–4^.

FAM171A2 is predominantly expressed in the brain, especially in hippocampal neurons^5^, the choroid plexus, in the vascular endothelium and microglia^6^. In addition, all three FAM171 family members have been identified at excitatory synapses in cultured neurons^7^ and FAM171A2 at hippocampal mossy fiber synapses^5^, suggesting a possible role at glutamatergic synapses. FAM171A2 variants are associated with neurodegenerative diseases, such as Alzheimer’s and Parkinson’s diseases, through the regulation of progranulin (PGRN) expression^6^. Whether FAM171A2 modulation of PGRN expression is direct or indirect remains unknown, and any potential FAM171A2 binding partners involved in this regulation are yet to be identified. FAM171A2 has been recently identified as a neuronal receptor mediating endocytosis of α-synuclein fibrils, and is thought to play a role in the spread of α-synuclein pathology in Parkinson’s disease^8^.

Structurally, all members of the FAM171 family of proteins are predicted to have an extracellular region of 282–321 residues in length, a single transmembrane domain, and larger intracellular domains spanning 452–566 residues. The predicted extracellular domain contains four putative N-linked glycosylation sites and whereas no intramolecular disulfide bonds are present across family members, FAM171B has a unique single unpaired cysteine in its extracellular domain. Overall sequence alignment shows the FAM171 proteins to have high sequence identity and similarity (**Supplementary Figure 1**), with FAM171A1 and FAM171A2 sharing greater sequence identity with each other than with FAM171B. Alignments of the predicted extracellular domains reveal high sequence conservation with FAM171A1 and FAM171A2, sharing a ∼45% sequence identity in their ectodomain. The most striking difference between the FAM171 sequences is a polyglutamine stretch of ∼14 residues at the N-terminus of FAM171B, that has been found to vary in length between 12 and 17 residues in humans^9^. Despite the high sequence identity, we will not discuss FAM171B in this work. There are currently no experimentally determined structures available for any FAM171 protein, and the SWISS-Model server^10^ fails to identify any structural homologs that cover more than 10% of each sequence.

In this study, we describe the three-dimensional structures of monomeric human FAM171A1 and FAM171A2 ectodomains, and we show that they both have a novel protein architecture. We demonstrate that the FAM171A1 and FAM171A2 ectodomains form equilateral trimers, with the propensity, for FAM171A1, to form larger trimer-trimer assemblies. Overall, our data show the quaternary structure of the ectodomain of the FAM171A1 and FAM171A2 proteins in solution, which suggest important functions in neuronal biology.

## RESULTS

### BIOPHYSICAL CHARACTERIZATION OF THE ECTODOMAINS OF FAM171A1 AND FAM171A2

To characterize the in-solution behavior of the extracellular domain (ectodomain) of human FAM171A1 and FAM171A2, we cloned their ectodomains, defined as the region between the first amino acid after the signal peptide and the last amino acid before the transmembrane domain (FAM171A1: T23-T303; FAM171A2: K30-T315), in frame with an IgG1-Fc fragment (**Figure 1A**).

**Figure 1:**
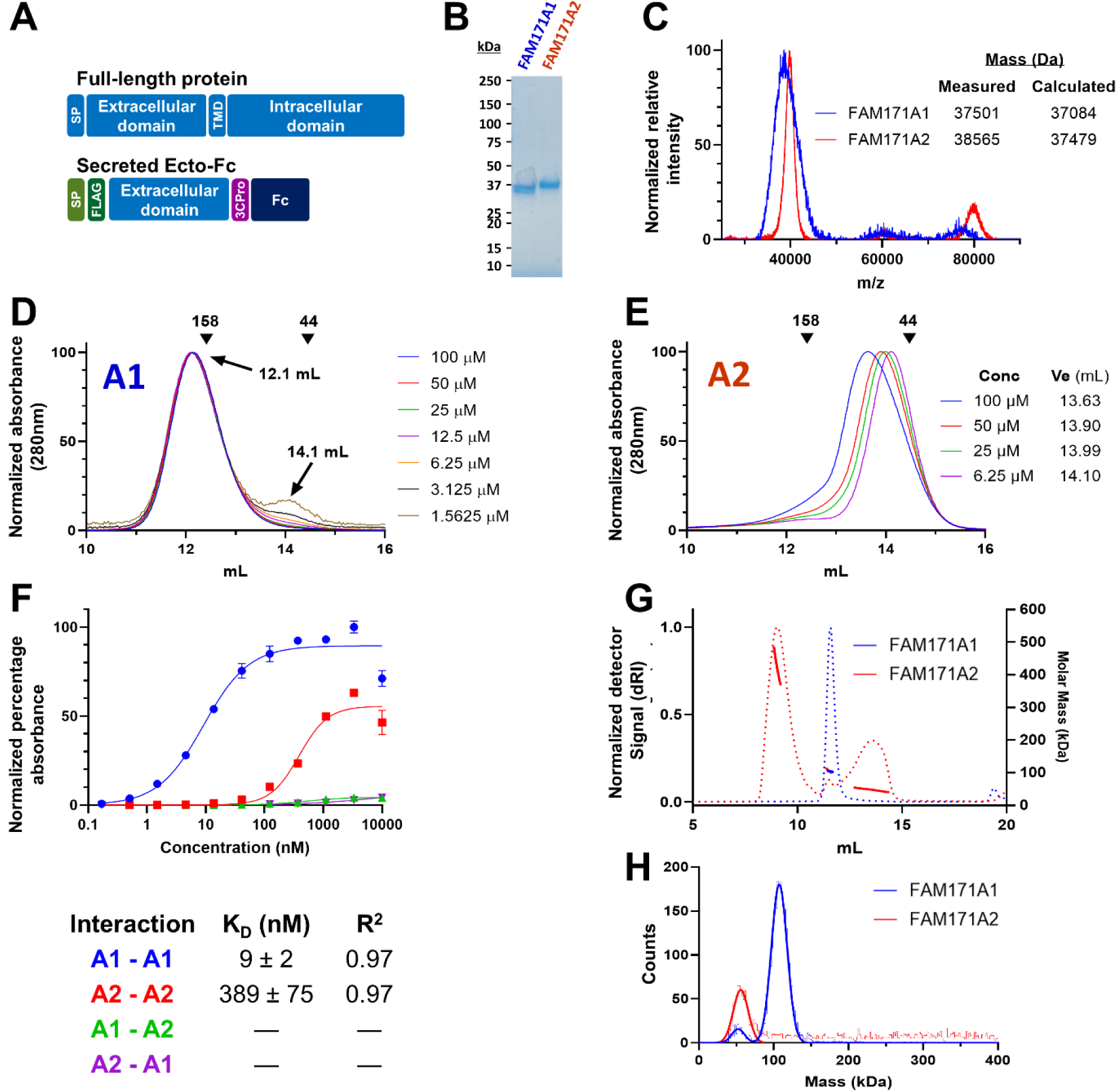
Biophysical characterization of FAM171A1 and FAM171A2. **A)** Schematic diagram of the domain organization of the FAM171 proteins. **Top:** architecture of the full-length protein, including the signal peptide (SP), and the transmembrane domain (TMD). **Bottom:** architecture of the Ecto-Fc protein, with the Prolactin signal peptide (light green, SP), the DYKDDDDK tag (dark green, FLAG), the HRV-3CPro cleavage site (magenta 3CPro) and the Fc- fragment (dark blue, Fc). **B)** Coomassie blue staining of the two FAM171 purified proteins separated by SDS-PAGE. **C)** MALDI-TOF of FAM171A1 and FAM171A2 showing the main peaks of the monomers. In the inset, the calculated MW for the amino acid sequence including four N-linked glycans. **D, E)** SEC profiles of FAM171A1 **(D)** and FAM171A2 **(E)** purified from HEK293S GnTI- cells at different concentrations, demonstrating oligomerization concentration-dependent. Both experiments were performed using a Superdex 200 10/300 GL column. Absorbance is normalized for comparison. Protein standard masses (kDa) indicated by arrowheads. **F) Top:** Affinity ELISA of FAM171A1 and FAM171A2 interactions. Purified Ecto-Fc FAM171 was titrated against Ecto-AP FAM171 proteins in an ELISA assay to assess relative binding affinities. Wells without AP-protein were used as a negative control to calibrate the zero response. **Bottom**: Table of the apparent affinities of the various combinations of AP-FAM171 and Fc-FAM171. Measurements were recorded in triplicate. **G)** SEC-MALS of FAM171A1 (blue) and Fam171A2 (red) demonstrating the difference in solution behavior. The graph shows the SEC results measured with a differential refractive index detector (dRI, dotted line trace, left Y axis), as well as protein mass distribution measured with multi-angle light scattering in parallel to dRI measurements (MALS, continuous line, right Y axis) represented as an overlay of both runs. Runs performed at 3 mg/ml (∼91µM) sample concentrations. Note: aggregation may appear in FAM171A2 over time, shown here at ∼9mL of elution. **H)** Mass photometry of FAM171A1 and FAM171A2. The graphs show an overlay of detected count numbers as a histogram and particle mass distributions represented by a Gaussian curve fitted to the main populations for FAM171A1 (blue) and FAM171A2 (red). FAM171A1 trimer remains stable even at concentrations as low as 10 nM, covering ∼90% of all detected molecules. The higher oligomeric states of the A2 ectodomain appear unstable at low concentrations, as indicated by the presence of only a monomer peak.

Transient expression in Expi293 cells and stable expression in HEK293S GnTI- cells (lacking the N-acetylglucosaminyltransferase I (GnTI) enzyme) were highly effective for all constructs, yielding >50 mg/L and >10 mg/L of purified protein, respectively, indicating that folding was not compromised by the cloning boundaries. Coomassie blue staining of the purified protein, and matrix-assisted laser desorption/ionization-time of flight mass spectrometry (MALDI-TOF MS) were used to measure the monomeric mass of FAM171A1 and FAM171A2 ectodomains purified from HEK293S GnTI- cells. As expected, masses for the ectodomain (FAM171A1, measured: 37.50 kDa, calculated 37.08 kDa; FAM171A2, measured: 38.56 kDa, calculated 37.48 kDa) (**Figure 1B, C**) were consistent with the calculated molecular weight (MW) based on the glycosylated amino acid sequence. These values include the N-terminal FLAG-tag and C-terminal cleaved 3CPro recognition site, and indicate these recombinant proteins are fully glycosylated with each of the four N-linked glycosylation sites occupied by a Man5GlcNAc2 moiety (1.23 kDa each), which are typical of GnTI- cells^11^.

Size exclusion chromatography (SEC) analysis of purified FAM171A1 showed a single peak at 12.1 mL with negligible change in elution volume across a 64-fold (100–1.562 µM, or ∼3,120–∼49 µg/mL) dilution series (**Figure 1D**). Column calibration demonstrates that this species is likely a stable oligomer larger than 160 kDa. At the lowest concentration measured (1.562 µM, or ∼49µg/mL), a small peak is present at 14.1 mL, suggesting a minor amount of oligomer dissociation. FAM171A2 shows a major multimeric species at 13.63 mL with a definite rightward shift in elution volume across a 16-fold dilution series (100–6.25 µM or ∼3,070–∼192 µg/mL), indicating that oligomer dissociation occurs at low protein concentrations (**Figure 1E**).

### THE ECTODOMAINS OF FAM171A1 AND FAM171A2 ARE OLIGOMERS IN SOLUTION

To calculate the molecular mass of each oligomer in solution, and by extension the stoichiometry of the ectodomain, we resorted to analytical ultracentrifugation (AUC). Sedimentation velocity data demonstrates that FAM171A1 at 20 µM (∼624 µg/mL) is a single species with a mass of ∼100 kDa, equivalent to 2.7 monomers (**Table 1, Supplementary Figure 2A**), assuming full N-linked glycosylation. FAM171A1 has an s_20,w_ = 5.35 S and a f/f_0_ = 1.53, suggesting an asymmetric structure. Sedimentation equilibrium experiments determine the mass of FAM171A1 to be 96.34 kDa, equivalent to 2.6 monomers, which is most consistent with a trimeric species. (**Supplementary Figure 3B**). In contrast, sedimentation velocity data under the same conditions indicate that FAM171A2 exists as two species (s_20,w_ = 3.24 S and s_20,w_ = 4.93 S) with an average f/f_0_ = 1.01, suggesting a smaller, more globular structure than FAM171A1 (**Supplementary Figure 3C**, **Table 1**). Consistent with SEC data, sedimentation equilibrium data best fits a FAM171A2 mixture consisting of 25% monomer (38.60 kDa) and 75% trimer (111.24 kDa). Overall, these biophysical results find FAM171A1 to exist as a single stable oligomeric species, likely a trimer, whereas FAM171A2 exists in a concentration-dependent monomer-trimer equilibrium.

**Table 1:** AUC analysis of FAM171A1 and FAM171A2 paralogues. The data and fits are shown in Supplementary Figure 3. Data were collected at 20 µM and 10 °C.

| Protein | Mass (kDa)<br>(Proportion, %) | | $\chi^2$ | $s_{20,W}$ (rmsd) | f/f<sub>0</sub> | Oligomer |
| --- | --- | --- | --- | --- | --- | --- |
|  | Species 1 | Species 2 |  |  |  |  |
| Sedimentation velocity experiments |  |  |  |  |  |  |
| FAM171A1 | ~100.0 (97) | - | - | 5.35 (0.002) | 1.53 | 2.7 |
| FAM171A1 | ND* (60) | ND (32) | - | 3.24, 4.93 (0.005) | 1.01 | - |
| Sedimentation equilibrium experiments |  |  |  |  |  |  |
| FAM171A1 | 96.347 (100) | - | 1.86 | - | - | 2.6 |
| FAM171A2 | 38.600 (25) | 111.239 (75) | 2.46 | - | - | 1, 2.9 |

\* Because this sample has multiple species with different frictional ratios, it is not possible to determine the mass using a single averaged frictional ratio.

To gauge the strength of the FAM171A1 and FAM171A2 oligomerization, an ELISA binding affinity assay was employed. The relative apparent affinity of each interaction was measured between AP-tagged FAM171 protein immobilized on the plate and the addition of a dilution series of purified Fc-tagged FAM171 proteins. Consistent with the SEC dilution series, the FAM171A1 self-association was ∼40-fold stronger (K_D_ = 9.0 ± 2.0 nM), than FAM171A2 self-interaction (K_D_ = 389 ± 74 nM), thereby explaining the rightward shift of the curve of FAM171A2 (**Figure 1F**). Despite their close sequence identity, no hetero-oligomerization (A1/A2) was detected. Overall, these experiments confirmed high-affinity FAM171A1 and FAM171A2 ectodomain homo-oligomerization, with no propensity for hetero-oligomerization, and further support their in-solution behavior.

We next used size-exclusion chromatography combined with multi-angle light scattering (SEC-MALS) and mass photometry to further investigate the oligomeric state of both FAM171A1 and FAM171A2 ectodomains. According to the MALS results for FAM171A1, the molecules in the single peak fraction from the SEC chromatogram had a mass of 104.3 kDa (± 0.1%), with monodispersity proven by the horizontal nature of the MALS curve. In contrast, the FAM171A2 sample resulted in three peaks, with the smallest fraction of the protein present in the trimeric form, followed by monomeric and higher molecular weight species. The molecules in the FAM171A2 sample showed polydispersity, likely due to polymerization or aggregation, which explains the sloped MALS curve at higher molecular weights (**Figure 1G**). Using mass photometry, purified FAM171A1 measured at 10 nM (∼0.31 µg/mL) showed a predominant trimer peak having a mean mass value of 107.3 kDa (σ 10.53 kDa), representing almost 90% of the detected molecules. Conversely, the FAM171A2 ectodomain sample displayed a wide range of particle masses, with the only small peak corresponding to the monomeric state (∼30% at 55.8 kDa, σ 9.97 kDa) (**Figure 1H**). SEC-MALS and mass photometry experiments support a stable trimeric state of FAM171A1 and a concentration-dependent monomer-trimer behavior for the FAM171A2 ectodomain.

### THREE-DIMENSIONAL STRUCTURE OF FAM171A1 ECTODOMAIN

To determine the cryo-EM structure of FAM171A1, grids were initially optimized using the FAM171A1 ectodomain protein at 10 mg/mL (∼312µM), supplemented with 0.02% NP-40. This initial experiment identified 20 nm round particles (**Figure 2A**), which enabled us to generate a three-dimensional map with a global resolution of 2.3 Å (**Figure 2B, C**). ModelAngelo^12^ was used for *de novo* fitting of a molecular model within this cryo-EM map, using the FAM171A1 extracellular domain amino acid sequence as an input. The initial model generated was a 60-mer, appearing as a hollow icosahedral sphere ∼220 Å in diameter, made up of 20 identical FAM171A1 trimers (**Figure 2D**). Each monomer has the N-terminus pointing to the surface of the sphere and the C-terminal end pointing inside the icosahedron. In the 60-mer, each FAM171A1 trimer is ∼70 Å across, and each monomer has three interaction interfaces, two with the two monomers completing the trimer, and a third interface in contact with one monomer of a neighboring trimer (**Figure 2E**). As we suspected that the 60-mer was the result of high protein concentration, new grids of FAM171A1 at 1 mg/mL (∼31 µM) and without NP-40 were imaged at a magnification of 81,000X. Data processing, particle selection, 2D classification, and 3D reconstruction using C3 symmetry, resulted in an even distribution of single trimeric particles of ∼70Å diameter (**Figure 2F-H**), identical to ones determined as components of the 60-mer. Together with the biophysical solution experiments, these data show that the ectodomain of FAM171A1 is a constitutive trimer. At a higher concentration and with a small amount of detergent, the isolated FAM171A1 ectodomain has a propensity to form an ordered hollow icosahedral sphere.

**Figure 2:**
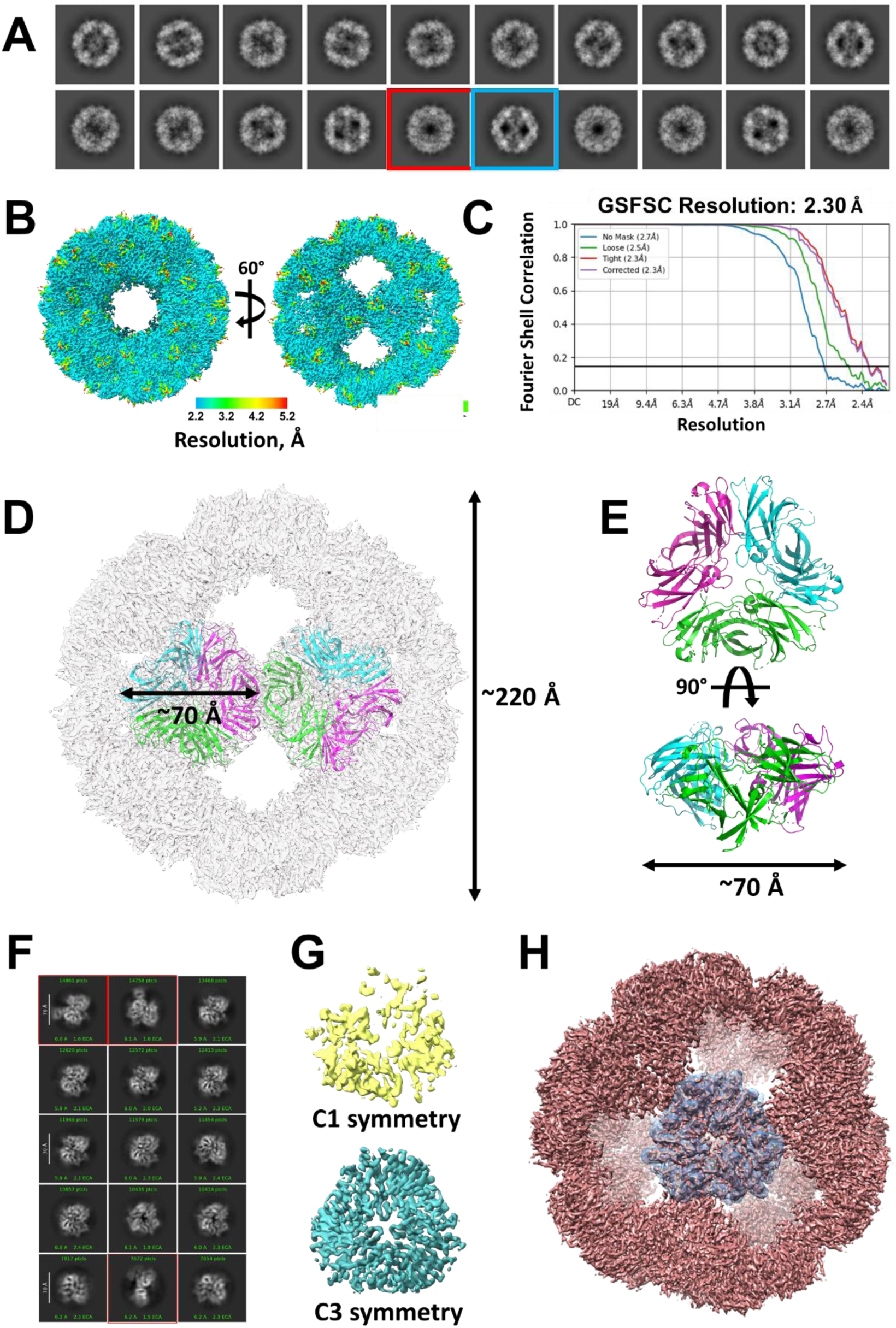
FAM171A1 cryo-EM maps. **A)** 2D class averages of FAM171A1 60-mer particles with the two main classes identified in different orientations. Protein concentration was 10 mg/mL (∼312µM). **B)** Local resolution map of the 60-mer particle with the maximum dimensions and two views differing of 60-degree rotation, similar to the views highlighted in **A**. **C)** Fourier Shell Correlation curve**. D)** Superimposition of two FAM171A1 trimers within the cryo-EM density. Measures detail the maximum dimension of the 60-mer (220 Å) and the trimer (70 Å). **E)** Refined FAM171A1 trimer model in two orientations. Each monomer has a different color. **F)** 2D class averages of FAM171A1 trimer. Protein concentration was 1 mg/mL (∼31µM). **G)** Three-dimensional reconstructions of the dataset shown in **F**, using C1 and C3 symmetry. **H)** Alignment of C3 symmetry FAM171A1 trimer map (gray) with FAM171A1 60-mer map (pink).

### THE STRUCTURE OF THE FAM171A1 MONOMER

The symmetry of the icosahedral sphere of FAM171A1 enabled us to achieve high-resolution of the structure of the constitutive monomer. Analysis of the three-dimensional structure of the monomeric ectodomain of FAM171A1 indicates that it is composed of two domains: residues T23-L118 form the N-terminal domain (Domain 1), which is followed by a tripeptide linker (P119-121) leading to the C-terminal domain (Domain 2), composed of residues S122 to P288 (**Figure 3A**). In the full-length protein, P288 is followed by a short sequence (I289 to P292) likely functioning as a stalk domain, leading to the transmembrane and cytoplasmic regions of the protein. Structural analysis of the FAM171A1 monomer using The Encyclopedia of Domains (TED) and CATH servers^13,14^ led to the identification of these two domains. Domain 1 is classified as 2.60.40.1120, a carboxypeptidase-like regulatory domain, and shows similarity to human carboxypeptidase D (PDB: 5AQ0) with structural alignment yielding an RMSD of 1.20 Å. In the CATH database, “2” is the class that comprises mostly beta sheets, “60” defines the architecture that comprises β-sandwich, “40” comprises Immunoglobulin-like domains, and “1120” defines the carboxypeptidase-like regulatory domain. Domain 2 is classified as 2.60.220, where “220” defines Chondroitinase proteins. Domain 2 belongs to the same immunoglobulin-like superfamily as the ZU5 domain, found in both human erythroid ankyrin and UNC5B. Despite this similarity, the ZU5 domain in the human erythroid ankyrin crystal structure (PDB: 3F59) aligns poorly with domain 2 of FAM171A1 (RMSD = 4.8 Å).

**Figure 3:**
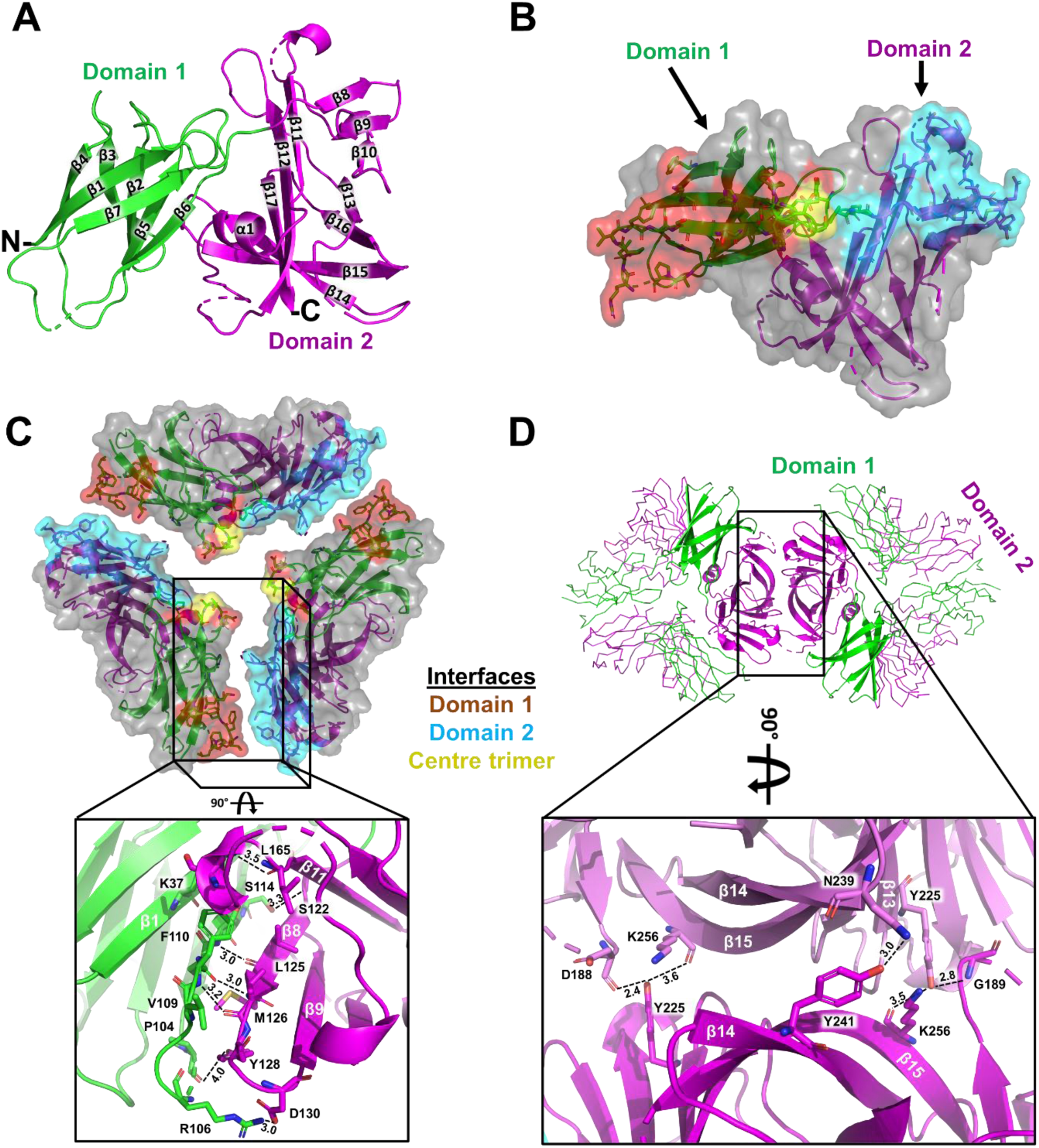
FAM171A1 ectodomain structure. **A)** FAM171A1 monomer structure with secondary structure elements labelled. Domain 1 (β1–7, green) and domain 2 (β8–17, magenta). N, N-terminus; C, C-terminus. Of note, the positions of residues 138-142 in the β10-β11 loop are unresolved, as well as residues 190-195 of the β11-β12, and residues 232-235 also remain unresolved. C-terminal residues 286-307 also remain unresolved in this structure. Domain 1 consists of β-sheets one and two, whereas domain 2 contains β-sheets three to five. **B)** FAM171A1 trimer surface contacts. Represented as a cartoon with a semi-transparent surface. Domain 1 interface, salmon; domain 2 interface, cyan; central interacting surface, yellow. Residues involved in the association are represented as sticks. **C)** Reciprocal FAM171A1 trimer interacting residues. Top-down representation of the FAM171A1 trimer with each monomer slightly separated for clarity. Inset highlights residues involved in trimer formation. Distances are in Å. **D)** FAM171A1 trimer-trimer interacting surface. Inset highlights residues involved in trimer-trimer formation. Distances are in Å.

Domain 1 is composed of two antiparallel β-sheets, which overall adopt a β sandwich structure (**Figure 3A**). Domain 2 is connected to domain 1 through the three-residue β7-β8 linker (**Figure 3A**). Overall, domain 2 adopts a more disordered β sandwich-like structure, consisting of two anti-parallel β-sheets and a six-residue α-helix. Interchain contacts are found between domain 1 of one monomer and domain 2 of a second monomer (**Figure 3B**). As searches with the SWISS-Model server^10^ failed to identify structural homologs covering more than ∼10% of each FAM171 sequence, to the best of our knowledge, this is the first structure where these two domains are combined, thus forming one novel overall structure.

### THE QUATERNARY TRIMERIC STRUCTURE OF THE FAM171A1 ECTODOMAIN

The most stable quaternary structure of FAM171A1 consists of three monomers forming an equilateral triangle (**Figure 2E, 3C**). Using the PDBePISA server^15^, buried and total surface areas were calculated for each monomer interaction as well as the whole trimer. Interactions between two monomers bury an area of ∼750 Å^2^ for each monomer. However, as each monomer interacts with two additional monomers, trimer formation buries a total area of ∼4,500 Å^2^ (∼15% of the total surface area). Residues involved in the formation of the FAM171A1 trimer are listed in **Supplementary Table 2**. The main contacts for trimer formation are made between β7 and β8 from each monomer, as well as loops between and flanking each of these β strands (**Figure 3C**). Hydrogen bonds between β1 and the β10-β11 loop also contribute to trimer formation. A salt bridge (D130-R106’) is also found between β6-β7 and β8-β9 loops at the outermost area of the interface. Contacts at the center of the trimer are also observed between the β1-β2 loop of each monomer. Analysis of the 60-mer structure indicates that the trimer-trimer interactions use most of the outer domain 2 surface opposite to the trimer center, with β13-15 strands of opposite monomers (**Figure 3D**) providing the key interactions (**Supplementary Table 3**). The four N-linked glycosylation sites of each monomer are all on superficial loops not implicated to interface formation.

### MUTAGENESIS OF FAM171A1 INTERFACES

As cryo-EM provided two quaternary FAM171A1 structures, a trimer and a 60-mer consisting of 20 equivalent trimers, mutants were designed to validate the observed oligomerization by disrupting the interfaces of these units, thus ruling out potential artifacts. Because of the extensive interactions at each interface, we reasoned that one or few Ala mutations may not be sufficient to disrupt these interfaces, and extensive and combinatorial Ala scanning may introduce further artifacts, including misfolding and endoplasmic reticulum retention^16^. Different FAM171A1 constructs were designed to introduce targeted N-linked glycosylation sites, creating a bulk that should prevent oligomerization (**Figure 4A, B**) without affecting folding or inducing protein aggregation. To block trimer formation, two mutations (F110N, S122N) were introduced, individually or together. Expression of all six mutants in Expi293F cells was successful, indicating that folding was not compromised by the introduction of additional N-linked glycosylation and aggregation was not observed for any of the FAM171A1 mutants. SDS-PAGE analysis showed each FAM171A1 mutant to be approximately 40 kDa in size, with mutants containing two introduced N-linked glycosylation sites migrating slightly slower than the single mutants, at approximately 45 kDa (**Figure 4C**). Proteins containing trimer-disrupting mutations (either F110N, S122N, or the double variant) were found to have smaller oligomerization compared to trimeric FAM171A1, eluting at 1.87 mL just before the 44 kDa standard on a Superdex 5/150 GL column, indicating that these targeted mutations disrupted trimerization (**Figure 4D**).

**Figure 4:**
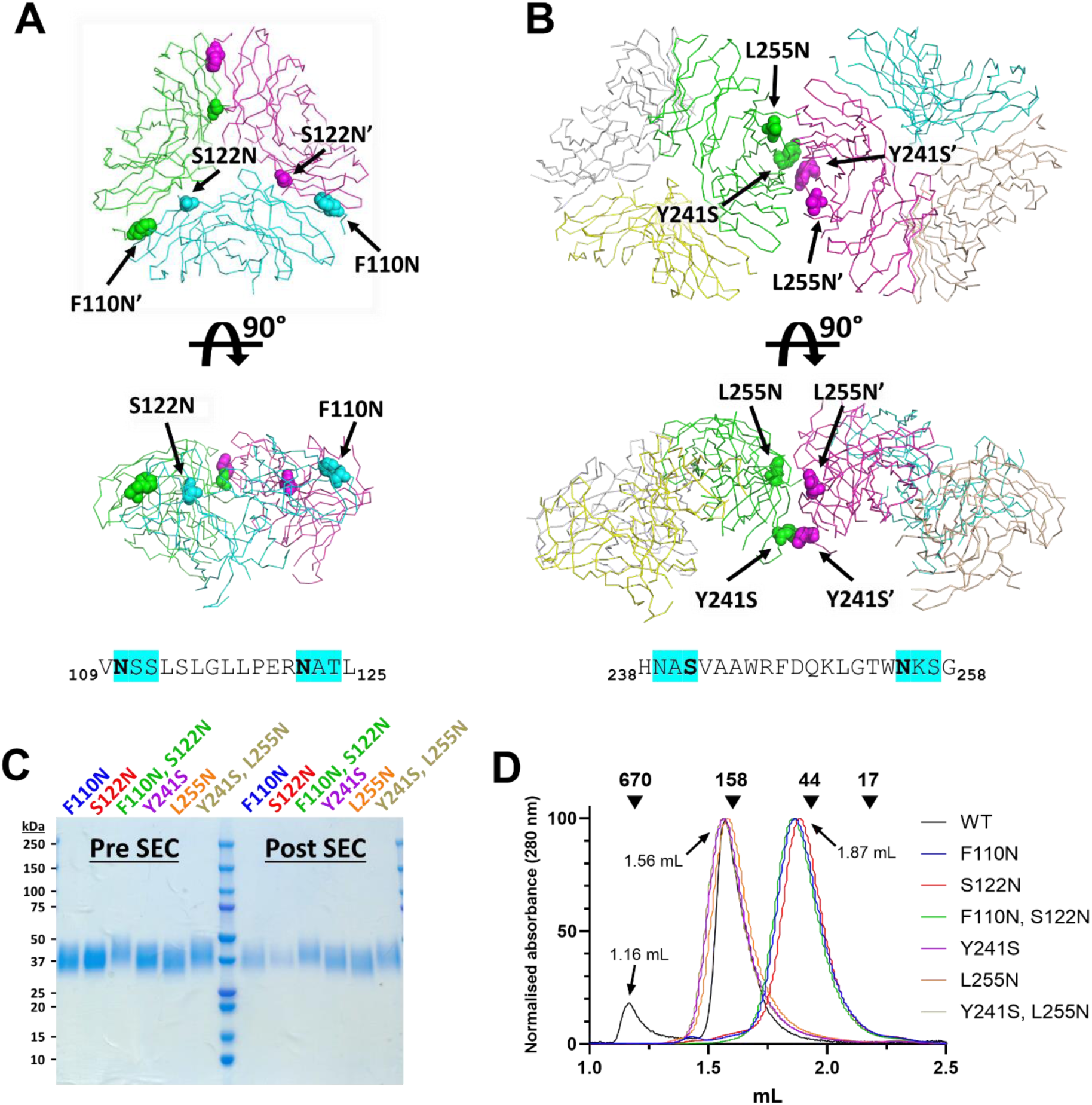
Mutagenesis of FAM171A1. **A-B)** FAM171A1 artificial N-linked glycosylation mutation sites. Trimer interface mutations S122N and F110N. Trimer-trimer interface mutations Y241S and L255N. FAM171A1 structures represented as ribbons with each monomer colored (**A**: green, cyan, magenta; **B**: yellow, grey, beige). Mutation sites represented as spheres. Bottom: amino acid sequences with the position of the new N-linked glycosylation sites. **C)** SDS-PAGE of purified FAM171A1 mutants before and after SEC injection and recovery. Note that the double mutants, having two additional N-linked glycosylation sites, migrate more slowly than the single mutants. **D)** SEC of FAM171A1 mutants. Normalized SEC analysis of purified FAM171A1 WT (Black) and mutants, using a Superdex 5/150 GL column. The group with a peak at 1.56mL (Y241S, L255N and Y241S/L255N) elutes as a trimer, but each mutant has lost the 60-mer form eluting at 1.16mL. The group that elutes at 1.87mL (S122N, F110N and S122N/F110N) elutes as a monomer. Mutants were all injected at ∼7.7mg/mL (∼240µM), except for S122N which was injected at ∼3mg/mL (∼93µM)

To confirm or rule out the presence of a FAM171A1 60-mer identified in cryo-EM analysis and SEC-SAXS experiments, we generated Y241S and L255N (and the double variant) mutations, which are located at the putative trimer-trimer interface. SEC showed the 60-mer disrupting mutants eluted at a volume comparable with wild-type trimeric FAM171A1 (1.56 mL). Taken together, the mutagenesis data show that both FAM171A1 trimer and trimer-trimer interactions generate quaternary structures and suggest that the icosahedral 60-mer is not an artifact of the conditions or additives used in cryo-EM or SAXS (see below) but can also be recapitulated using size-exclusion chromatography only.

### THE TRIMERIC STRUCTURE OF THE FAM171A2 ECTODOMAIN

The purified ectodomain of FAM171A2 (amino acids P36-M293) was used for determining a high-resolution structure of FAM171A2 by cryo-EM. Initial imaging conditions for FAM171A2 were similar to the ones used for FAM171A1, 10 mg/mL (∼307µM) supplemented with 0.02% NP-40. These conditions provided well-distributed and randomly oriented particles suitable for three-dimensional reconstruction. The FAM171A2 map generated when applying C3 symmetry appeared to be reliable when assessing the local resolution map and a global resolution of 2.78 Å indicated the map to be of an overall high quality (**Figure 5A-C**). The refined trimeric FAM171A2 structure largely overlaps with the FAM171A1 trimer structure, with all-atom alignment in PyMOL^17^ providing an RMSD of 1.49 Å. Unresolved residues are located in β9-β10, β10-β11, β11-β12, and β13-β14 loops, each missing two to five amino acids (residues 147–148, 165–166, 197–201 and 240–241, respectively). Structural analysis of the FAM171A2 monomer shows that domain 2 of FAM171A2 has a slightly worse alignment with the ZU5 domain (RMSD = 5.1 Å) than the FAM171A1 ortholog. Interactions between each FAM171A2 monomer bury an area of approximately 790 Å^2^, making the total buried surface upon trimer formation ∼4750 Å^2^. Contacting residues in the FAM171A2 trimer are listed in **Supplementary Table 4**. Similar to FAM171A1, the major contacts for trimer formation are made between β7 and β8, as well as the loops between and on either side of these β strands (**Figure 5D, E**). The β10- β11 loop has hydrogen bonds with β1 and the β6-β7 loop also, as does β6 with the β9-β10 loop. Similar to FAM171A1, contacts at the center of the trimer involve the β1-β2 loop of each monomer. Overall, the three-dimensional trimeric structures of FAM171A1 and FAM171A2 trimers are nearly identical, as expected from their high sequence similarity, with both showing similar interfaces for trimer formation. One notable difference, however, giving the identical imaging condition between FAM171A1 and FAM171A2 (∼10 mg/mL and 0.02% NP-40), was the absence of the 60-mer particle in the FAM171A2 grids. This observation further strengthens the idea that the icosahedral form of FAM171A1 is not an artifact of the experimental conditions and we speculate that it likely has some biological relevance.

**Figure 5:**
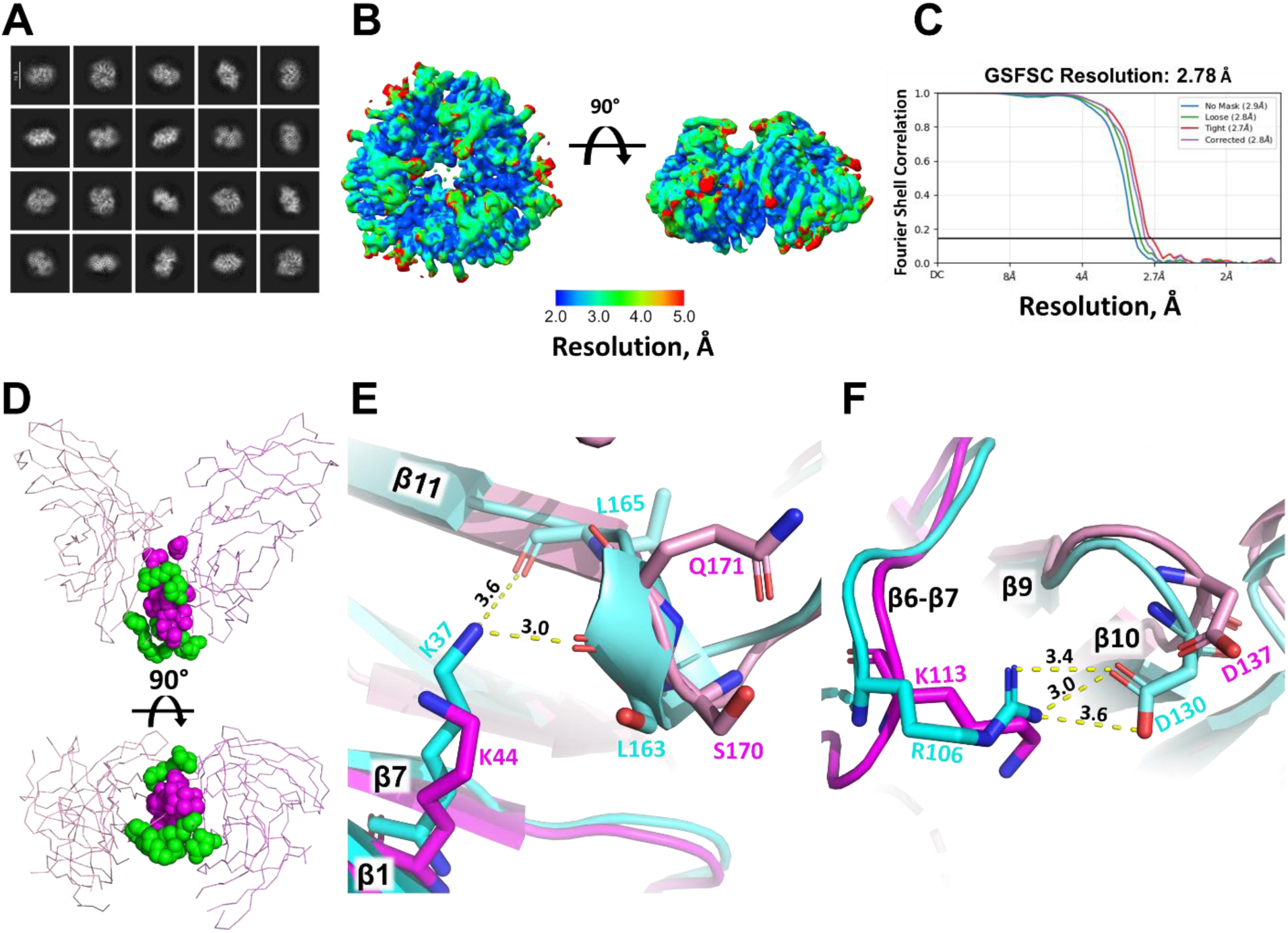
FAM171A2 ectodomain structure and comparison with FAM171A1. **A)** 2D class averages of FAM171A2 trimer particles. **B)** Local resolution map. **C)** Fourier Shell Correlation curve. **D-F)** Comparison of FAM171A1 and FAM171A2 trimer-forming residues. **D)** FAM171A1 model, showing two monomers in shades of pink involved in trimer formation. Hydrogen bonding residues shared with FAM171A2 are highlighted as magenta spheres whereas FAM171A1-specific interactions are highlighted as green spheres. **E)** FAM171A1 (cyan) K37-L163 and L165 hydrogen bonds and equivalent FAM171A2 (pink) residues. **F)** FAM171A1 (cyan) P104-Y128 hydrogen bond and R106-D130 salt bridge and equivalent FAM171A2 (pink) residues. Interacting residues are shown as sticks and colored according to chain. Bond distances (Å) highlighted by yellow lines. Secondary structure elements labelled in white, residues colored according to chain. Dashed yellow lines depict polar interactions with atomic distance given in Å. Heteroatom colors: oxygen (red), nitrogen (blue). Shades of cyan and pink indicate different monomers.

### SMALL ANGLE X-RAY SCATTERING (SAXS)

To validate the quaternary assembly in solution of FAM171A1 and FAM171A2 solved by Cryo-EM, we used SAXS, to accurately measure particle dimensions in solution. Data were collected using a SEC-SAXS with a co-flow setup at the Australian Synchrotron. Purified FAM171A1 and FAM171A2 samples were used at concentrations similar to the ones used in Cryo-EM (3-11.4 mg/mL or ∼94 to 355µM), in a Tris buffer of physiological ionic strength (**Figure 6A**). P_(r)_ plots for FAM171A1 and FAM171A2 have a bell shape (**Figure 6B**), with minimal tailing at high r values, indicative of a globular protein shape. The calculated maximum dimension (D_max_) of each protein is larger than what was calculated by Cryo-EM, with FAM171A1 (132 Å) and FAM171A2 (125 Å), likely because the envelope includes the flexible N-linked glycans, as well as the flexible N- and C- termini, which are absent in the cryo-EM reconstructions. For all SAXS samples, Guinier plots are linear (**Figure 6C**) as expected for non-aggregating monodisperse samples^18^. The Guinier analysis indicated that FAM171A1 has a radius of gyration (Rg) of 37.9 Å ± 0.4 Å, slightly larger than FAM171A2 (36.8 ± 0.3 Å) (**Table 2**). The raw data plot of the FAM171A1 sample at 11.4 mg/mL has at least three distinct centers of scattering and is strikingly different from the decay profiles of the smaller FAM171 species. With an Rg of 101.9 ± 0.1 Å and D_max_ of 249 Å the SAXS data are consistent with the 60-mer species. The solution scattering measurements (**Table 2**) demonstrate that these protein preparations are suitable for further structural modelling. DAMMIF^19^ and DENSS^20^ modelling programs were used to generate low-resolution models from each scattering dataset, using both P1 and P3 symmetries. DAMMIF and DENSS provided similar results, with ***χ***^2^ values (0.207–0.248). Multiple runs of data modelling for FAM171A1 and FAM171A2 trimers were performed, with each atomic model fitting the triangular density model with excellent correlation scores (0.705 and 0.696, respectively). Multiple models of the data of FAM171A1 60-mer model also fitted the icosahedral sphere density well, with a correlation score among all the different runs of 0.777 (**Figure 6D**). Taken together, the SAXS data supports the FAM171A1 and FAM171A2 trimer structures, as well as the icosahedral structure of FAM171A1. This is particularly notable as none of the SAXS samples contained NP-40 detergent, and for the 60-mer structure, the protein concentration was 11.4 mg/mL (∼355µM), thus similar to that used in cryo-EM.

**Figure 6:**
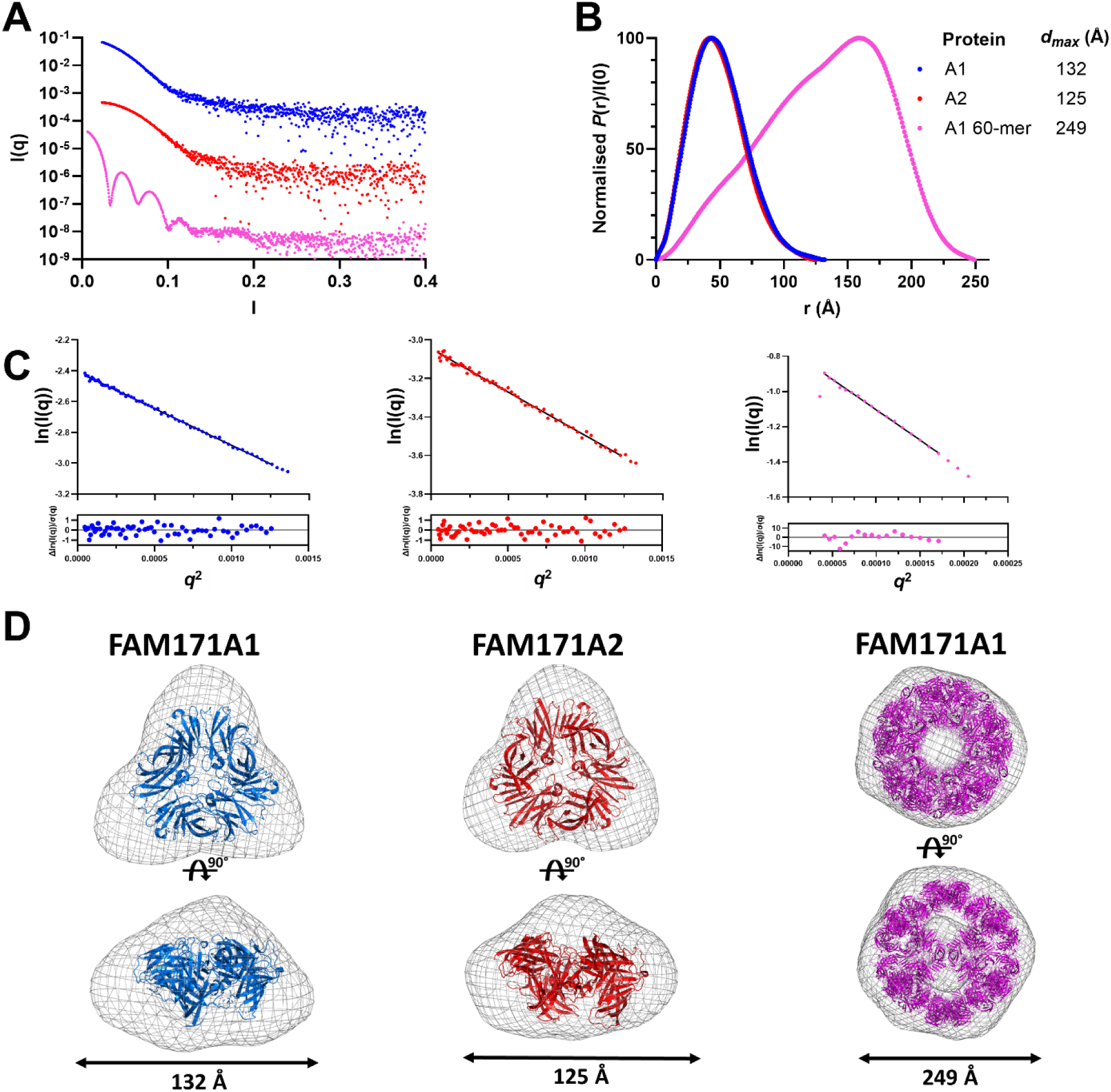
Small angle X-ray scattering data of FAM171A1 and FAM171A2 ectodomains. **A)** Scattering decay profiles (offset on the Y axis for clarity) of the highest concentrations of FAM171 proteins, color coded as in B. **B)** Normalized P_(r)_ functions of FAM171 proteins indicate the distribution of all interatomic distances and the maximum dimension (D*_max_*) of the particles, in Å. See also Table 2 for a complete report of other SAXS parameters. **C)** Guinier analyses where the solid black line represents the linear fit through the Guinier region. Below the linear Guinier, the scattering of the fit shows excellent linearity of the Guinier region. **D)** Envelope reconstruction of the scattering data overlaid to the refined cryo-EM structures.

**Table 2:** FAM171 SAXS data analysis and modelling.

| Protein | FAM171A1 | FAM171A1<br>(60-mer) | FAM171A2 |
| --- | --- | --- | --- |
| Beamline | SAXS/WAXS | BioSAXS | SAXS/WAXS |
| **Guinier analysis (RAW)** |  |  |  |
| $I(0) \pm \sigma$ (cm<sup>-1</sup>) | 0.0900 ± 0.0003 | 0.4689 ± 0.0004 | 0.0475 ± 0.0003 |
| $R_g \pm \sigma$ (Å) | 37.9 ± 0.4 | 101.9 ± 0.1 | 36.8 ± 0.3 |
| $qR_g$ range<br>(datapoint range) | 0.2580 – 1.3435<br>(0 – 61) | 0.6545 – 1.3315<br>(1 – 17) | 0.2505 – 1.2872<br>(0 – 60) |
| Linear fit assessment, $R^2$<br>(AUTORG fidelity) | 0.9983 | 0.9981 | 0.9938 |
| **$P(r)$ analysis (RAW)** |  |  |  |
| $I(0) \pm \sigma$ (cm<sup>-1</sup>) | 0.0899 ± 0.0007 | 0.4654 ± 0.0001 | 0.0476 ± 0.0003 |
| $R_g \pm \sigma$ (Å) | 38.9 ± 0.3 | 98.90 ± 0.01 | 36.8 ± 0.3 |
| $D_{\max}$ (Å) | 132 | 249 | 125 |
| $q$ range (Å<sup>-1</sup>) | 0.0233 – 0.6406 | 0.0064 – 0.4208 | 0.0068 – 0.6406 |
| $P(r)$ reciprocal-space fit: C<br>$\chi^2$ | 0.2431 | 2.3651 | 0.2479 |
| **Volume estimates (Å<sup>3</sup>)** |  |  |  |
| Porod volume (Å<sup>3</sup>) (ratio<br>$V_P$/calculated $M$) | 171,200 | 2,360,000 | 160,300 |
| **Molecular mass $M$**<br>**estimates (Da)** |  |  |  |
| From chemical composition | 32,144 | 32,144 | 32,539 |
| From c-independent<br>method<br>(Bayesian inference, range<br>with % confidence) | 134,300 – 162,700<br>(97%) | 1,120,000 – ∞<br>(98%) | 127,500 –<br>151,400<br>(93%) |

## DISCUSSION

FAM171 is a family of three type I cell surface receptors sharing high sequence identity within the family and predicted structural similarities. Considering their association with numerous human diseases, ranging from multiple forms of cancer metastasis^2–4,21–23^ to neurodegenerative diseases, such as Alzheimer’s and Parkinson’s diseases^6,8^, they have the potential to drive many different context-dependent signaling pathways and cellular responses. Despite these associations, little information is available on their biochemical, biophysical, and cell biological features.

In this work, we provide new insights into the molecular properties of FAM171A1 and FAM171A2. First, we show that the monomeric ectodomain of human FAM171A1 and FAM171A2 has a novel architecture generated by a combination of two domains. Second, their ectodomains oligomerize as homo-trimers, which in the case of FAM171A1, have also the propensity to form larger trimer-trimer assemblies at high concentrations.

Using cryo-EM, we solved the monomeric structures of FAM171A1 and FAM171A2, which is a novel combination of two known domains, classified in known CATH superfamilies as reported in the Encyclopedia of Domains (TED). Interestingly, the overall architecture was correctly predicted by AlphaFold as deposited in the Protein Structure Database^24^. The discovery of this novel architecture is significant because it suggests that the extracellular domain of the FAM171 family of proteins may have evolved through the duplication of a carboxypeptidase-like regulatory domain and the ZU5 domain. Moreover, the availability of these new structures may facilitate the identification of other proteins with uncharacterized but related architectures.

Domain 2 (residues S122 to P288) is reported in TED by encompassing residues 121–307. With the experimental structure at hand, it appears that residues P289–V294 lay outside the globular portion of domain 2, likely forming a short stalk domain that connects to residues T295–L307, which are predicted to participate in the transmembrane domain and appear as an alpha-helix in the TED-predicted structure.

The structure of the FAM171A1 and FAM171A2 ectodomain showed that their monomers oligomerize to form a stable homo-trimer in solution, and likely on the cell surface, as the three C-termini leading to the three transmembrane domains are oriented in the same direction (e.g. toward the cell membrane). The differences in the affinity and stability of FAM171A1 versus FAM171A2 trimers likely originate from the additional stabilizing interactions unique to the FAM171A1 interface. In FAM171A1, two hydrogen bonds (K37-L163’ and K37-L165’) and a salt bridge (R106-D130’), located above and at the outer periphery of the shared interactions, appear to strengthen the shared hydrogen bonds between anti-parallel β strands. These FAM171A1-specific interactions likely account for the enhanced stability of the FAM171A1, which remains a trimer over a large concentration span (from ∼312µM to ∼10nM). By ELISA, we ascertained that FAM171A1 and FAM171A2 do not interact with each other to form a hetero-oligomer, therefore their endogenous ligand(s) remain unknown. We speculate that trimerization may facilitate ligand-receptor clustering on the cell surface, leading to intracellular signals possibly linked to actin stress fiber formation and cytoskeletal organization, which are essential for cell shape and motility^1^. Alternatively, it is possible that trimerization is stable only in the absence of an endogenous ligand; when a high-affinity ligand is present, the trimer may dissociate to form the hetero-complex in a 1:1 or other quaternary stoichiometries. To further investigate the role of individual monomer-monomer interactions, alanine scanning of residues within the trimer interface, followed by SEC analysis of the resulting proteins, could provide insights into the contributions of specific hydrogen bonds and salt bridges to FAM171A1 trimer stability. However, given the extensive interface interactions, multiple mutations may impair folding and expression, therefore impairing this type of study. FAM171A2 levels are increased in the cerebrospinal fluid of Parkinson’s disease patients and FAM171A2 promotes the spread of alpha-synuclein pathology in mouse models^8^. FAM171A2’s domain 1 was recently shown to bind alpha-synuclein and promote the spread of alpha-synuclein pathology in mouse models^8^. The high-affinity interaction involves Domain 1 of FAM171A2 and α-synuclein preformed fibrils. It will be important to establish experimentally how the trimeric structure of both FAM171A1 and FAM171A2 interact with α-synuclein preformed fibrils. Interestingly, the trimeric quaternary structure of FAM171A1 has the propensity to form larger trimer-trimer assemblies (**Figure 2D**) using the curved β13-15 strands of domain 2 of opposite monomers (**Figure 3D**). *In vitro*, the 60-mer hollow icosahedral sphere of FAM171A1 (but not FAM171A2) forms at a high protein concentration (∼10mg/mL or ∼312µM), and was observed in cryo-EM (**Figure 2A**), SEC (**Figure 4D**), and SEC-SAXS experiments (**Figure 6B)**, in the presence or absence of detergent NP-40, indicating this is not an artifact of a specific additive or experimental setup. Moreover, either Y241S or L255N mutants completely abolished the formation of the 60-mer, strengthening the idea that the extended assembly is biologically relevant. However, the icosahedral sphere is only energetically favored in solution because each ectodomain lacks the transmembrane and intracellular domains, both of which would be oriented toward the center of the icosahedron, rendering this assembly incompatible with cell membrane tethering. However, one could theoretically flatten the 60-mer into a two-dimensional array of proteins, with all transmembrane domains pointing toward the cell surface.

Furthermore, a weak trimer-trimer interaction, compared to the stability of individual trimers, could enable FAM171A1 to be present in multiple oligomeric states on the cell surface, allowing for reversible receptor clustering. While further conjecturing is possible, more research in cells and *in vivo* is needed to elucidate the role of oligomerization.

Determining the physiological and pathophysiological roles of FAM171A1 and FAM171A2, the protein domains involved, and trimer or monomer configurations underlying these functions, will be an important focus of future research. Additionally, identification of endogenous ligands for FAM171A1 and FAM171A2 will be essential to elucidating their specific biological roles.

### LIMITATION OF THE STUDY

This study was conducted *in vitro*, using highly purified protein, encoding only the ectodomain of either protein. Thus, we have no structural information on the entire protein. We discussed the biological relevance of the trimer and the 60-mer, however, no cell biological experiments were performed to identify these structures and a potential two-dimensional array on the surface of the cell.

## METHODS

### Gene synthesis of FAM171A1 and FAM171A2 ectodomains

Amino acid sequences were derived from the UNIPROT database (Uniprot.org) for human Fam171A1 (Q5VUB5) and Fam171A2 (A8MVW0). The cDNA sequence of each ectodomain was custom synthesized by Gene Universal (Newark, DE, USA) and cloned in frame as C-terminal Fc or AP-fusion proteins using 5’ NotI and 3’ XbaI sites into modified pCMV6-XL4 expression vectors. Between the ectodomain of the protein of interest and the Fc domain, there is a 3CPro cleavage site (LEVLFQ/GP) that was used to remove the Fc for purification purposes. The AP (alkaline phosphatase) construct differs from the Fc construct as it lacks the 3CPro cleavage site, contains AP in place of the Fc and carries a His6 tag at its C-terminus, in frame with the AP. The fact that AP is enzymatically active provides a simple way to test expression and measure accurate protein concentration. Both Ecto-Fc and Ecto-AP carry an N-terminal DYKDDDDK tag inserted after the prolactin leader peptide.

### HEK293T-17 transfection

HEK293 T-17 human embryonic kidney cells were obtained from American Type Culture Collection (ATCC), cat# CRL-11268. The cells were grown in Dulbecco’s modified Eagle’s medium (DMEM; Invitrogen) supplemented with 10% fetal bovine serum (FBS; Invitrogen) and penicillin/streptomycin (Invitrogen). One day before transfections, cells were plated on 6-well culture plates (Greiner) at equal densities. Cells were transfected with 1μg of either FAM171A1 or FAM171A2 plasmids using polyethylenimine (Polysciences). One day after transfection, cells were lysed with SDS loading buffer and boiled at 90°C for 5 min.

### Protein Expression

Stable cell lines expressing each FAM171 Fc-tagged construct were generated using adherent HEK293S GnTI- cells. HEK293S GnTI- cells were obtained from American Type Culture Collection (ATCC cat# CRL-3022). These cells lack N-acetylglucosaminyltransferase I (GnTI) activity, and consequently glycosylation remains restricted to a homogeneous seven-residue oligosaccharide^11^. These cells were grown in Dulbecco’s Modified Eagle Medium (DMEM) supplemented with 5% fetal bovine serum (FBS), maintained in a humidified incubator at 5% CO2 and 95% air. Stable cell line generation using the Fc vector (pCMV6-XL4) requires co-transfection with an empty pcDNA3.1 to confer geneticin (G418) resistance, at a ratio of 3:1 (pCMV6-XL4:pcDNA3.1). Cells were transfected using the calcium phosphate method, followed by G418 selection (500 µg/mL) (Sigma)^25^. Resistant clones were isolated using Pyrex cloning rings (Corning) and their protein expression was assessed via western blot analysis. The best expressing clones were amplified, frozen and used for large-scale protein expression, which was achieved in Nunc TripleFlask cell culture flasks until the appropriate amount of medium was collected. When expressed transiently, FAM171A1 or FAM171A2 plasmids were transfected in Expi293™ cells (Thermo) using ExpiFectamine™ (Thermo) according to the manufacturer’s instructions. Conditioned media was harvested four days after transfection and again two days after replenishing with fresh medium.

### SDS-PAGE

Samples were boiled at 95 °C for 5 min and subsequently loaded on a 4-20% polyacrylamide gels. Separated proteins were then stained with SimplyBlue Coomassie stain for 1h at RT and detailed overnight in water.

### Protein purification

Proteins were affinity purified using CaptivA™ PriMAB rProtein A Affinity Resin (Repligen) at a rate of 1–2 mL/min by gravity flow. Saturated resin was washed extensively (50mM Tris HCl, pH 8.0, 450mM NaCl) and equilibrated (50 mM Tris HCl, pH 8.0, 150 mM NaCl, 1 mM DTT). The Fc fragment was cleaved with HRC-3C protease (10µg/mL of enzyme in 50 mM Tris HCl, pH 8.0, 150 mM NaCl, 1 mM DTT) and incubated overnight. Eluted protein was concentrated to 2-10 mL with Vivaspin concentrators (Sartorius-Stedim) and further purified by size exclusion chromatography using a HiLoad 16/600 Superdex 200 PG (Cytiva, Marlborough, MA, USA). For analytical size exclusion chromatography, 100 µL samples were loaded onto a Superdex 200 10/300 GL column (Cytiva) equilibrated in 20 mM Tris HCl, pH 8.0, 150 mM NaCl. Protein was used immediately or aliquoted and flash-frozen in liquid nitrogen and stored at -80C until needed.

### ELISA relative binding affinity assay

A 50 µL solution of 3 µg/mL of mouse anti-AP (IgGAb-1 clone 8B6.18; Thermo) in PBS was added to each well of a 96-well plate and incubated at 4°C overnight. The following day, plates were washed three times with PBS-T (0.01% Tween-20 in PBS) and blocking buffer (5% non-fat milk in PBS) was added to each well for one hour. Subsequently, blocking buffer was removed and 40 µL conditioned media containing Ecto-AP was added to each well at room temperature. The plates were washed once with PBS-T and 40 µL of purified Ecto-Fc at different concentrations (∼10,000 to ∼17nM) were added to each well and incubated for three hours at room temperature. The plates were washed one time with PBS-T and 40 µL of a solution of 1.25 µg/mL monoclonal mouse anti-human IgG1-HRP (BioRad) in 1% non-fat milk in PBS was added to each well and incubated for one hour at room temperature. Plates were then washed three times with PBS-T buffer, and 50 µL 1-Step Ultra TMB-ELISA HRP substrate (Thermo) was added to each well. The absorbance at 650 nm was measured using EnSpire 2300 Multilabel Reader (Perkin Elmer) every five minutes for one hour. Data was baseline corrected, normalized and analyzed in GraphPad Prism 10 using the specific binding with Hill slope least squares fit function.

### SEC-MALS

The SEC-MALS setup consists of a Superdex 200 Increase 10/300 GL gel filtration column (Cytiva) connected to a 1260 Infinity II HPLC system (Agilent), an inline 1260 Infinity II VWD UV detector (Agilent), an Optilab T-rEX refractive index detector (Wyatt Technology), and a DAWN 8+ multi-angle light scattering detector (Wyatt Technology). Protein samples were prepared at 3 mg/ml (∼93µM) in a buffer containing 20 mM Tris pH 7.4, 150 mM NaCl. A standard curve was generated using bovine serum albumin (BSA) at 1 mg/ml in the same buffer condition. Molar mass calculations were based on angular Rayleigh scattering data and a known protein-specific refractive index increment (dn/dc) of 0.185 ml/g. The molecular weights of the peak content were calculated in ASTRA software (Wyatt Technology, version 7.3.2). The resulting data were exported to GraphPad Prism 10 for visualization and interpretation.

### Mass photometry

Mass photometry experiments were performed using a Refeyn OneMP instrument with the droplet-dilution method in normal acquisition mode and standard image size. The sample chambers were prepared by attaching silicon gaskets (Grace Biolabs) to clean High Precision cover slips (Marienfeld). The correct position for data acquisition after focusing was determined by the empty buffer signal and was accepted at values below 0.7. Molecular mass distributions were derived from a calibration curve generated the same day from a triplet of measurements for NativeMark Unstained Protein Standard (Invitrogen) diluted in sample buffer (20 mM Tris pH 7.4, 150 mM NaCl), with a maximum error of 3.76%. Working protein concentrations were estimated by running samples of 10, 15, 30, and 50 nM dilutions in the sample buffer, finding a concentration giving total count values between 2,000-3,000. Final data were collected at protein concentrations of 10 nM (∼3.1µg/mL) for A1 and 30 nM (∼9.6µg/mL) for A2, with each measurement performed at least in triplicate. The data were then analyzed using Discover MP software (Refeyn) and exported as calibrated molecular mass values. For every measurement, histograms were generated using frequency distribution analysis with a mass bin width of 1.9 kDa in GraphPad Prism 10. The histogram data were then analyzed using a Gaussian nonlinear regression model.

### MALDI-TOF mass spectrometry

A 5800 TOF/TOF MALDI mass spectrometer (Sciex) was used to analyze the samples. The mass spectrometer was equipped with a Nd:YAG laser operating at 355 nm. The instrument was regularly calibrated using ProteoMass™ Protein MALDI-MS Calibration Kit (Sigma). The instrument was controlled by TOF/TOF Series Explorer software^26^ (Sciex). It was operated in linear mode with the laser intensity was optimized for each run, and 400 laser shots per spot were collected. At least three separate spots per protein were shot and one representative dataset was chosen for analysis.

### Analytical ultracentrifugation

Sedimentation velocity and equilibrium experiments were performed using either a Beckman Coulter Optima or XLI analytical ultracentrifuge with an An-50Ti rotor pre-cooled to 10°C. Protein samples were prepared to a concentration of 20 µM (∼640 µg/mL) in a buffer of 20 mM Tris, pH 8.0, 150 mM NaCl, and this same buffer was used in the reference cell. All experiments were conducted at 10 °C to minimize aggregation, since the equilibrium experiments were conducted over four days. Solvent density, solvent viscosity and estimates of partial specific volumes for each protein (including post-translational modifications) were calculated using SEDNTERP^27^. For the sedimentation velocity experiments, 380 µL of sample and 400 µL of reference buffer were loaded into 1.2 cm double-sector cells with quartz windows. Samples were centrifuged at 50,000 rpm and radial absorbance data was collected at 295 nm. Sedimentation velocity data were fit using SEDFIT^28^, including RI and RA noise correction. For the sedimentation equilibrium experiments, 110 µL of sample and 120 µL of reference buffer were loaded into 1.2 cm double-sector cells with quartz windows. Samples were centrifuged sequentially at 8,000, 10,000, 14,500 and 17,500 rpm for at least 24 hours at each speed. The radial absorbance data collected at 293–297 nm at each 24-hour time point. Sedimentation equilibrium data were fit using SEDPHAT^29^, fitting for the bottom and baseline but without any noise correction, and including two scans for each speed taken ∼1 hour apart demonstrate equilibrium had been reached.

### Cryo-EM sample preparation and data collection

Purified FAM171A1 and FAM171A2 proteins were prepared to a concentration of ∼10 mg/mL (∼310µM). Holey carbon grids (R 1.2/1.3 300 Mesh Quantifoil) were glow-discharged for 30 seconds at 15 mA using GloQube (Quorum) and 4 µL of protein was spotted on each grid. NP-40 (Sigma-Aldrich) at 0.02% was added to the protein solution right before applying the sample on the grid, blotted for 4 seconds with a blot force of -1 using a Vitrobot Mark IV (Thermo Fisher Scientific) set to 22°C and 100% humidity and plunge frozen into liquid ethane. The FAM171A1 sample without detergent was prepared in the same way as described above, except that the protein concentration was reduced to 1 mg/mL (∼31µM).

The samples were imaged on a Titan Krios G4 electron microscope (Thermo Fisher Scientific), equipped with a K3 direct electron detector and BioQuantum imaging filter (Gatan), operating at 300 kV in counting mode with a 10 eV energy filter slit. Cryo-EM data collection was performed using the automated software package, EPU v3.6 (Thermo Fisher Scientific). Movie stacks were recorded at a nominal magnification of 81,000 for FAM171A1 plus NP-40 and 105,000 for FAM171A1 without NP-40 and FAM171A2 plus NP-40 with calibrated pixel sizes of 1.069 and 0.833 Å/pixel, respectively. The dose rate was 15 e^−^/pixel/s, yielding a total electron dose of 50 e^−^/Å^2^, fractioned into 50 frames. In total, 5,593 micrographs were collected for FAM171A1 plus NP-40, 4,509 for FAM171A1 without NP-40 and 6,100 for FAM171A2 plus NP-40. Cryo-EM data acquisition statistics are shown in **Supplementary Table 1.**

### Cryo-EM data processing

All cryo-EM data processing was conducted using CryoSPARC v4.5.0^30^. Beam-induced motion was corrected, and movie frames were aligned using Patch Motion Correction and Contrast Transfer Function (CTF) estimation was performed using Patch CTF estimation. To pick particles for the FAM171A1 and FAM171A2 trimer, the template was generated from a small subset of the denoised micrographs by using reference-free automated particle picking, Blob Picker, followed by particle extraction and 2D classification. The selected 2D classes were used as a template to pick particles using Template Picker. The particles from the FAM171A1 60-mer dataset were picked using Blob Picker, with the ring blob option enabled. Particle picking was inspected using the Inspect Picks tool. 300,479 particles for the FAM171A1 60-mer, 1,529,470 particles for FAMA171A1, and 2,140,563 particles for FAM171A2 trimer were extracted. The extracted particles were subjected to iterative rounds of 2D classification (See **Supplementary Figure 2** for details). The particles in 2D classes were selected to generate the 3D reconstruction. The initial model was obtained *ab initio* and used for subsequent refinement. The final maps were obtained after rounds of non-uniform and heterogenous refinement followed by CTF refinement and Reference Based Motion Correction. The refined particles generated the final maps at 2.30 Å (FAM171A1 60-mer, Icosahedral symmetry), 3.35 Å (FAM171A1 trimer, C3 symmetry) and 2.78 Å (FAM171A2 trimer, C3 symmetry). The global resolutions were calculated according to the gold standard FSC criterion of 0.143.

### Model building

Initial placement of an AlphaFold 3 generated FAM171A1 trimer in the 60-mer map used the ‘dock in map’ function in Phenix^31^. For comparison, a *de novo* model was also generated using Model Angelo^12^. Coot was used for manual refinement of the AlphaFold generated model. The manually refined model was then subject to real-space refinement in Phenix. The model was subject to iterative cycles of manual and real space refinement before validation using the ‘comprehensive validation’ function in Phenix.

### Small angle X-ray scattering (SAXS)

SAXS data collection: SAXS experiments were carried out at the Australian Synchrotron using the SAXS/WAXS or BioSAXS beamlines, equipped with a Pilatus3 2M and Pilatus3X 2M detector, respectively (both with pixel sizes of 172 x 172 µm). Both beamlines have a photon energy of 12.0 keV (1.0322 Å wavelength). The sample to detector distance was set to 2700 mm, providing an s range of approximately 0.005–0.27 Å. Purified protein samples (3–10 mg/mL; ∼94-312µM) in a buffer of 20 mM Tris, pH 8.0, 150 mM NaCl, were prepared in a 96-well plate. 50 µL of protein was automatically loaded into the system, subject to on-line SEC using a Superdex 200 Increase 5/150 GL (Cytiva) column at 0.2 mL/min, before passing through the capillary co-flow system. Scattering data was collected once per second for the duration of the run.

#### SAXS data analysis

- All SAXS data was processed and analyzed using ATSAS^32^ and RAW^33^ software packages. Scattered intensity was plotted against s. Each sample was checked for non-linearity in the low s region, an increase in intensity indicating aggregation. RAW was used to calculate linear Guinier plots for *s* · *R_g_* < 1.3. GNOM^34^ was used for indirect Fourier transform, providing P(r) which defines the relative probability of scattering centers and the maximum dimension of the particle (*D_max_*). CRYSOL^35^ was used to generate theoretical scattering curves from cryo-EM atomic coordinates, and was used to compare against experimental scattering curves. DAMMIF^19^ and DENSS^33^ were used to generate *ab initio* models from the scattering data and SUPALM^36^ used to superimpose *ab initio* models onto AlphaFold 3 and cryo-EM models.

## AUTHOR CONTRIBUTIONS

Tom W Bird led the construct design, protein expression, purification, SEC characterization, cryo-EM of FAM171A2, MALDI-TOF, ELISA, mutagenesis, structure refinement, and structure analysis. Sepideh Valimehr led the cryo-EM of FAM171A1 and cryo-EM data refinement for FAM171A1 and A2; Gerd Mittelstädt and Florian De Pol contributed to the initial protein purification and characterization; David Wood and Zachary Tillet led the AUC experiments; Leanid Kresik led the SEC-MALS and mass photometry experiments; Davide Comoletti, Eric Hanssen, Joris de Wit, Renwick CJ Dobson, Dimphna Meijer, Tom W Bird designed the study, analyzed the data, and wrote the manuscript. All authors have contributed to the manuscript editing.

## RESOURCE AVAILABILITY

Requests for further information and resources and reagents should be directed to and will be fulfilled by the lead contact, Davide Comoletti

## DATA AND CODE AVAILABILITY

All cryo-EM maps and atomic models have been deposited to the Electron Microscopy Data Bank and Protein Data Bank. All EMDB and PDB accession codes are listed in the key resources table. They are publicly available as of the date of publication.

This paper does not report original code.

Any additional information required to re-analyze the data reported in this paper is available from the lead contact upon request.

## ACKNOWLEDGMENTS

This work was funded, in part, by National Science Foundation, award 1755189, to D.C.; by the Wellington Medical Research Foundation, 2019/301, and the School of Biological Sciences at Victoria Scholarship to TB. This work was supported by the following funding to R.C.J.D: 1) the New Zealand Royal Society Marsden Fund (contract UOC1506); 2) a Ministry of Business, Innovation and Employment Smart Ideas grant (contract UOCX1706); and 3) the Biomolecular Interaction Centre (University of Canterbury). J.D.W. was supported by funding from FWO project grants G0C4518N, G0A8720N and G0A8320N; Queen Elisabeth Medical Foundation for Neurosciences; Stichting Alzheimer Onderzoek-Fondation Recherche Alzheimer (SAO-FRA) standard grant 20230028, and a KU Leuven Methusalem grant. We would like to sincerely thank Dr. Christine Orengo (University College London) for helpful advice and discussion on the CATH database.

---

**Supp Figure 1:**
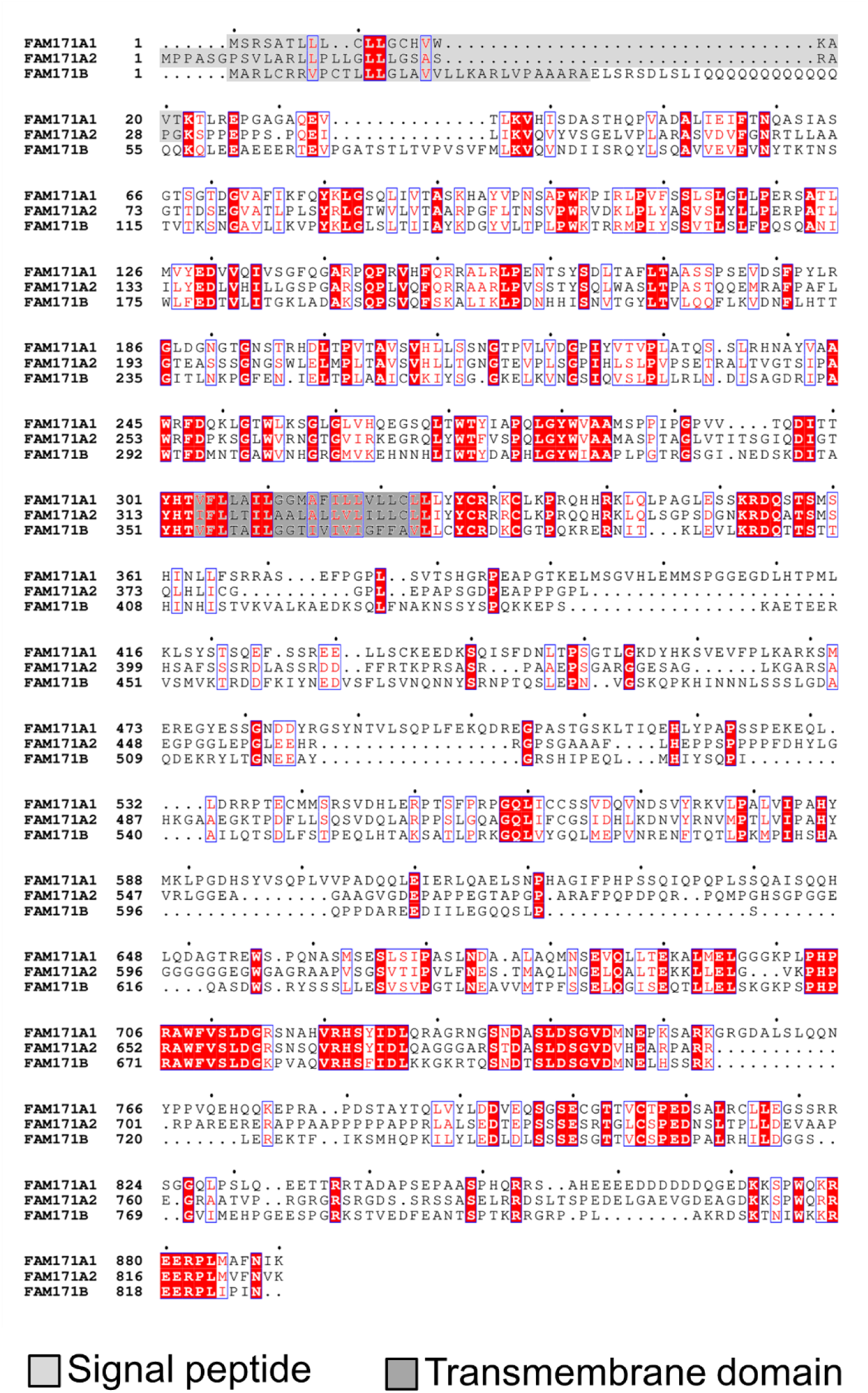
FAM171 sequence alignment. Multiple sequence alignment using Clustal Omega. Leader peptide (light grey) and transmembrane domains (dark grey). Conserved residues (identity) in all three paralogs have a red background, similar residues have a blue box around. Gaps in sequence are signaled by a dot.

**Supp Figure 2:**
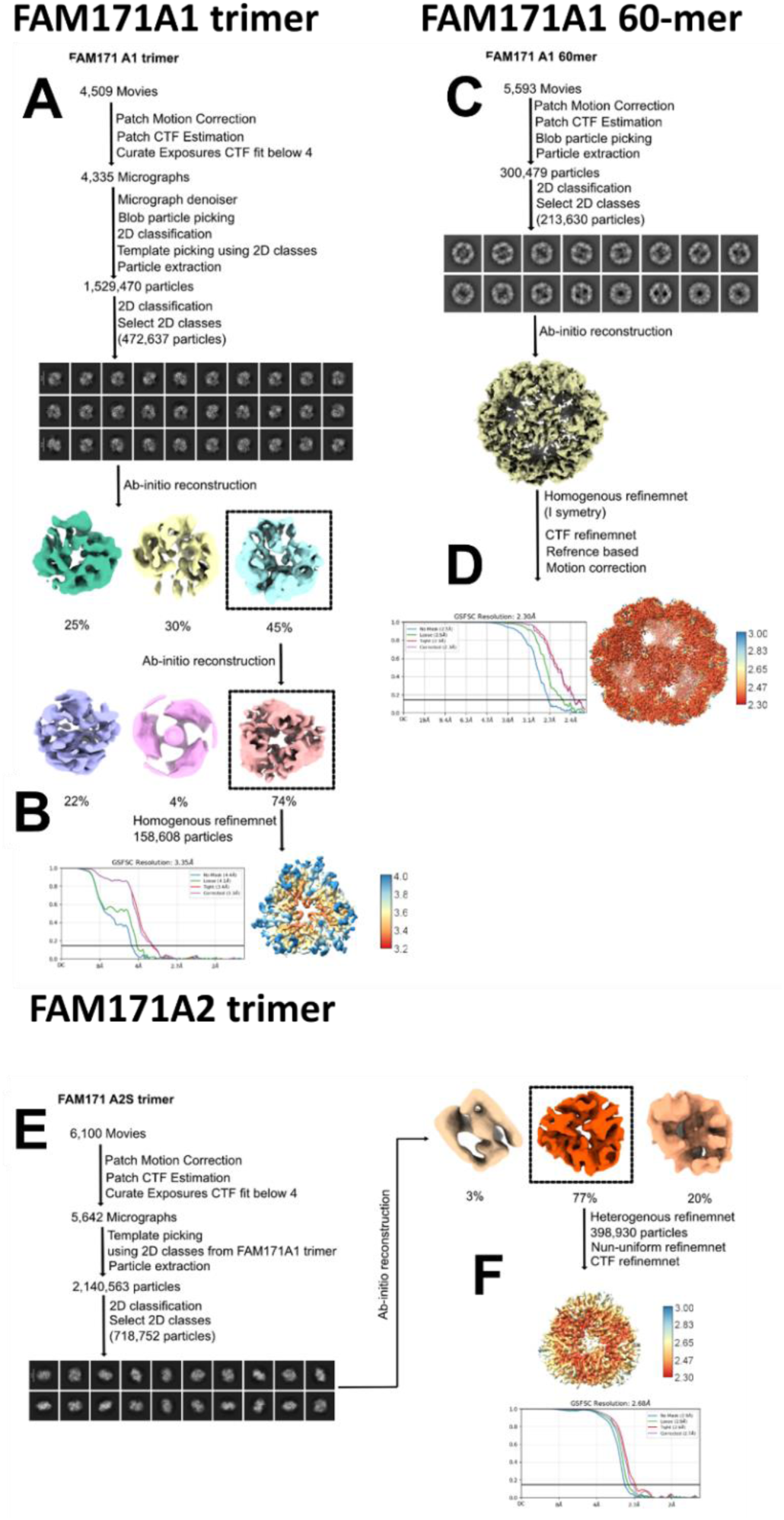
Cryo-EM data processing workflow for FAM171A1 and FAM171A2 trimers and FAM171A1 60mer. **A)** Cryo-EM data processing workflow and selected 2D classification to obtain 3D reconstruction of FAM171A1-trimer (left) and FAM171A1-60mer (right). **B)** Final 3D reconstruction and Fourier Shell correlation (FSC) curve of FAM171A1-trimer (left) and FAM171A1-60mer (right). **C)** Cryo-EM data processing workflow for FAM171A2S trimer. Cryo-EM data processing workflow and selected 2D classification to obtain 3D reconstruction of FAM171A2S-trimer. **D)** Final 3D reconstruction and Fourier Shell correlation (FSC) curve of FAM171A2S-trimer.

**Supp Figure 3:**
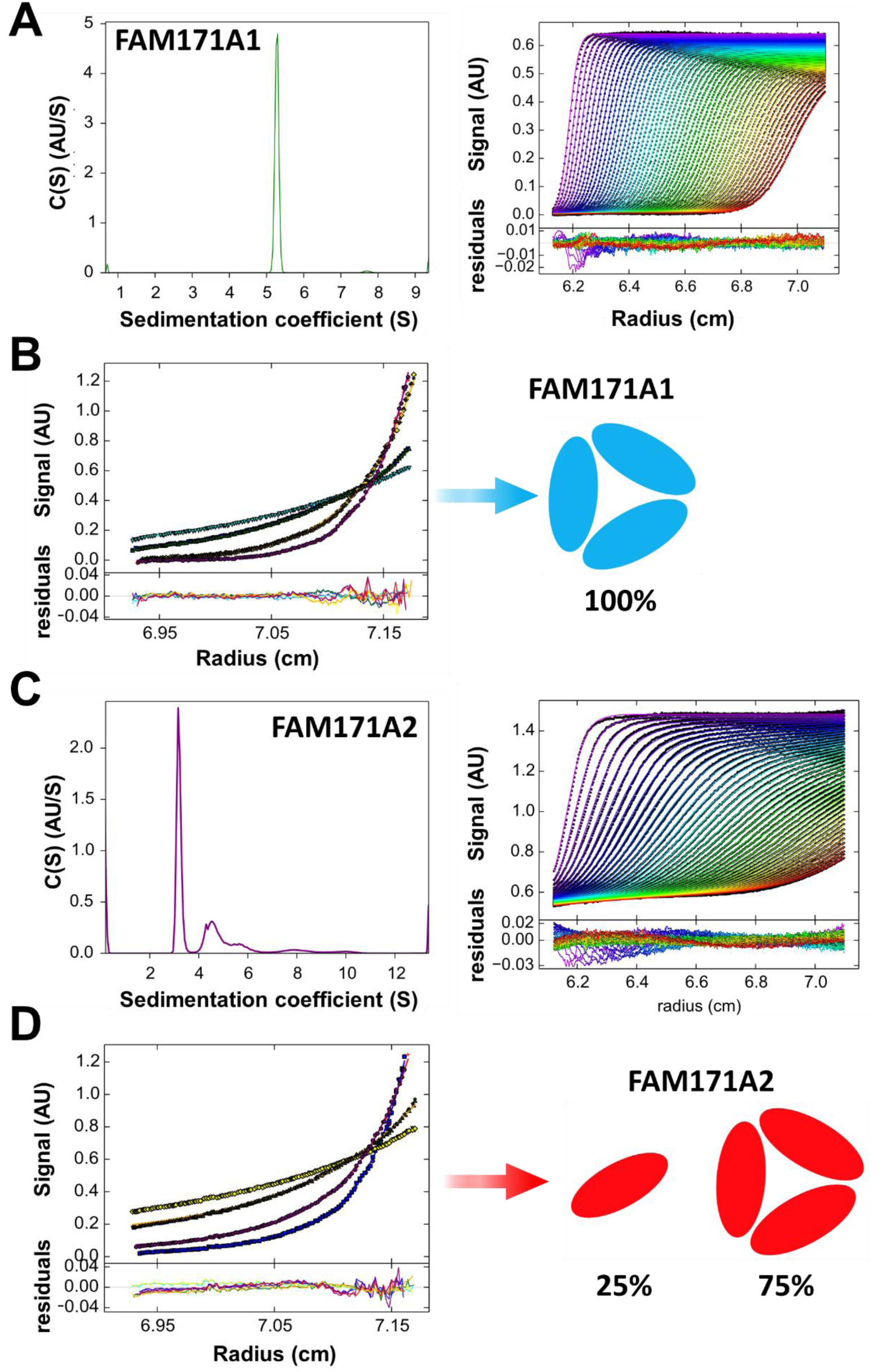
Analytical ultracentrifugation of FAM171A1 and A2 ectodomains. The sedimentation velocity experiments (**A, C**) were conducted at a concentration of 20 μM (∼624µM), at 10 °C and 50,000 rpm. The sedimentation equilibrium experiments (**B, D**) were in the same buffer and protein concentration, but at 8,000, 10,000, 14,500 and 17,500 rpm. Equilibrium data were fit globally using SEDPHAT to determine molecular masses. **A)** Continuous size distribution for FAM171A1 on the left showing the presence of a single species. Raw data and fits with residuals are shown on the right (rmsd = 0.002). Data were fit using SEDFIT to determine the sedimentation coefficients (s_20,w_ = 5.35) and frictional ratio (f/f_0_ = 1.53). For clarity, every 4^th^ scan and 3^rd^ data point are shown. **B)** Sedimentation equilibrium analysis of FAM171A1 fitted to a single discrete component model (global reduced *χ*2 = 1.86), giving a mass of 96.347 kDa, most consistent with a trimer species (a schematic of the oligomerization of the result is on the right). **C)** Continuous size distribution for FAM171A2 on the left showing the presence of a single species. Raw data and fits with residuals are shown on the right (rmsd = 0.005). Data were fit using SEDFIT to determine the sedimentation coefficients (s_20,w_ = 3.24, s_20,w_ = 4.93) and frictional ratio (f/f_0_ = 1.01). For clarity, every 10^th^ scan and 3^rd^ data point are shown. **D)** Sedimentation equilibrium analysis of FAM171A2 fitted to a two discrete component model (global reduced *χ*2 = 2.46), giving mass of 38.600 kDa (fixed) and 111.239 kDa (fitted), most consistent with a monomer-trimer self-association (a schematic of the oligomerization of the result is on the right). Fits to monomer-tetramer and dimer-trimer models result in a worse fit.

**Supp Figure 4:**
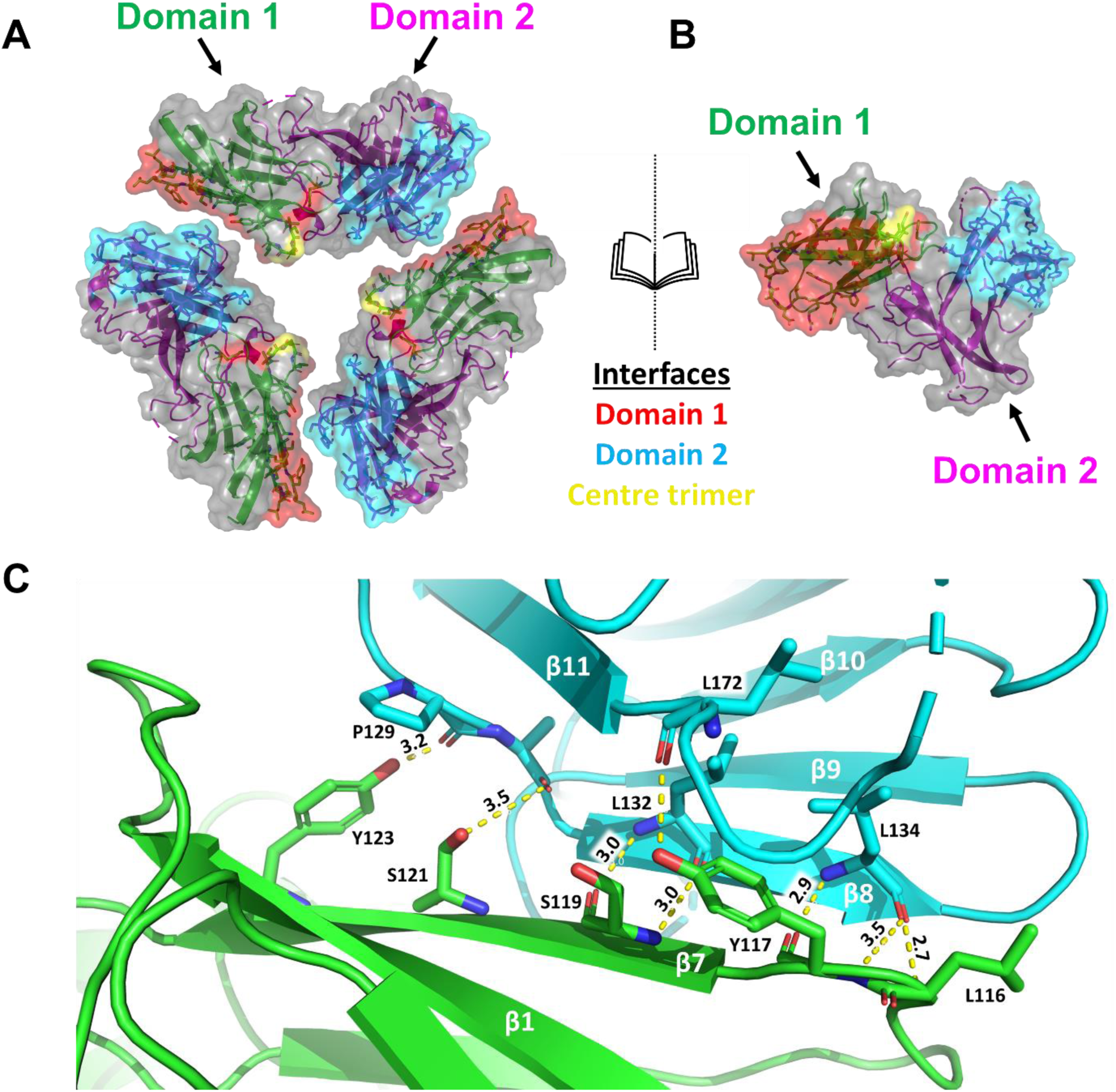
FAM171A2 surface contacts and key trimer interface residues. **A)** Exploded view of the FAM171A2 trimer represented as a cartoon with a semi-transparent surface. Domains 1 and 2 are colored green and magenta, respectively. Residues involved in the association are represented as sticks. **B)** Side-on representation of one monomer. **C)** Zoomed-in view of the interface of two FAM171A2 monomers (green and cyan) with interacting residues shown as sticks and colored according to chain. Bond distances (in Å) are shown as yellow lines. Secondary structure elements labelled in white, residues labelled in black. Heteroatom colors: oxygen (red), nitrogen (blue).

**Supplementary Table1:** Cryo-EM data collection, refinement, and validation statistics.

|  | **FAM171A1-60mer** | **FAM171A1-trimer** | **FAM171A2-trimer** |
| --- | --- | --- | --- |
| **Data collection and processing** |  |  |  |
| Magnification | 81,000 | 105,000 | 105,000 |
| Voltage (kV) | 300 | 300 | 300 |
| Electron exposure (e<sup>-</sup>/Å<sup>2</sup>) | 50 | 50 | 50 |
| Defocus range (μm) | 0.5-1 | 0.6-1.2 | 0.6-1.2 |
| Pixel size (Å) | 1.069 | 0.833 | 0.833 |
| Symmetry imposed | I | C3 | C3 |
| Initial particle images (no.) | 300,479 | 1,529,470 | 2,140,563 |
| Final particle images (no.) | 213,630 | 158,608 | 398,930 |
| Map resolution (Å) | 2.30 | 3.35 | 2.68 |
| FSC threshold | 0.143 | 0.143 | 0.143 |
| **Refinement** |  |  |  |
| Initial model used | AlphaFold | FAM171A1-60mer | AlphaFold |
| Model resolution (Å) | 2.3 |  | 2.8 |
| FSC threshold | 0.5 |  | 0.5 |
| Model resolution range (Å) |  |  |  |
| Model composition |  |  |  |
| Non-hydrogen atoms | 5179 |  | 5383 |
| Protein residues | 687 |  | 735 |
| *B*-factors (Å<sup>2</sup>) |  |  |  |
| Protein (mean) | 30.75 |  | 128.03 |
| R.m.s. deviations |  |  |  |
| Bond lengths (Å) | 0.004 |  | 0.006 |
| Bond angles (°) | 0.643 |  | 0.623 |
| Validation |  |  |  |
| MolProbity score | 1.26 |  | 1.74 |
| Clashscore | 4.89 |  | 9.54 |
| CaBLAM outliers (%) | 0.17 |  | 1.56 |
| Rotamer outliers (%) | 0.54 |  | 0.74 |
| Ramachandran plot |  |  |  |
| Favoured (%) | 99.53 |  | 96.44 |
| Allowed (%) | 0.47 |  | 3.56 |
| Disallowed (%) | 0 |  | 0 |
| PDB ID | 9Y63 |  | 9Y6N |
| EMDB ID | 72539 |  | 72623 |

**Supplementary Table 2.**
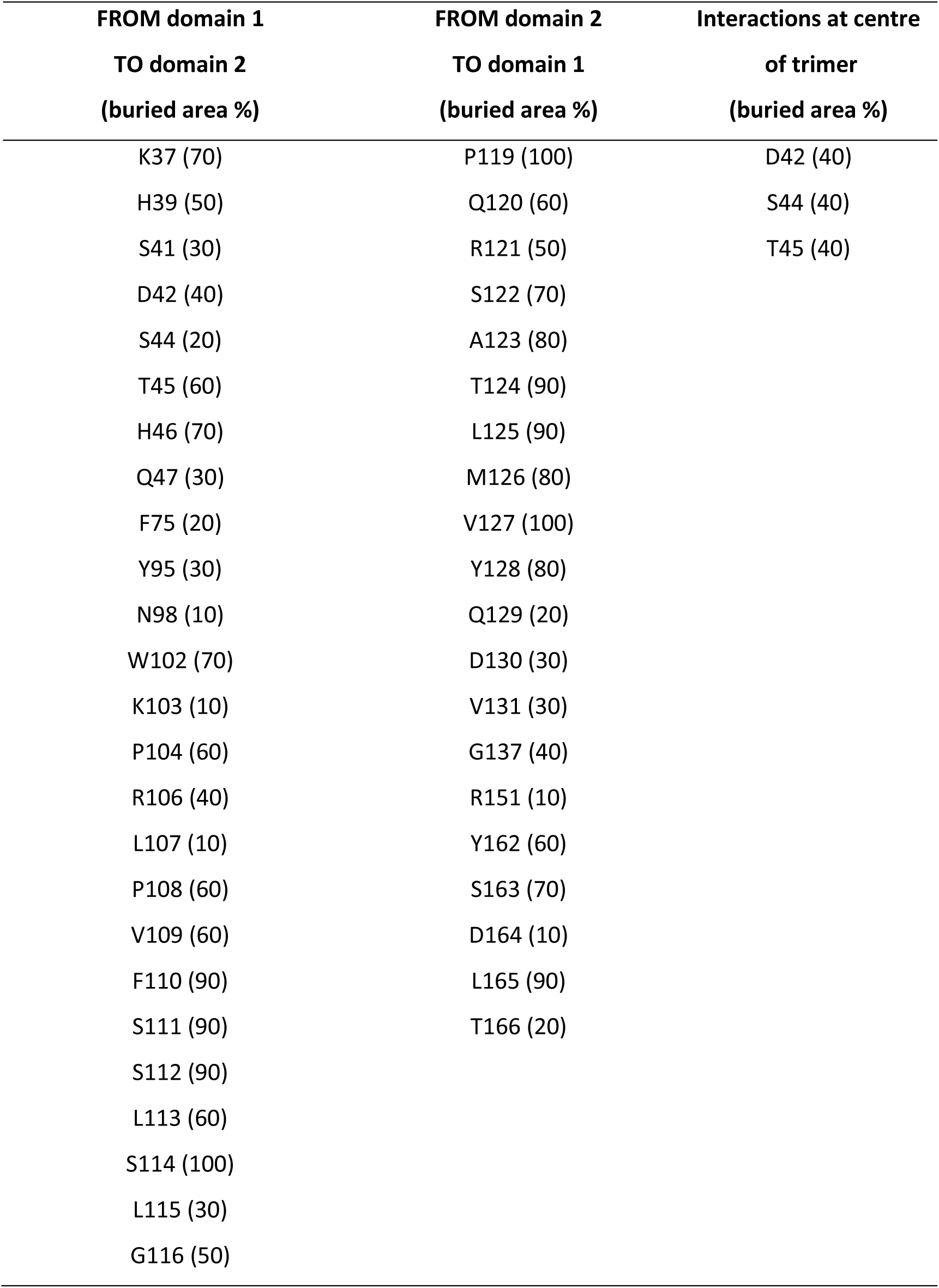
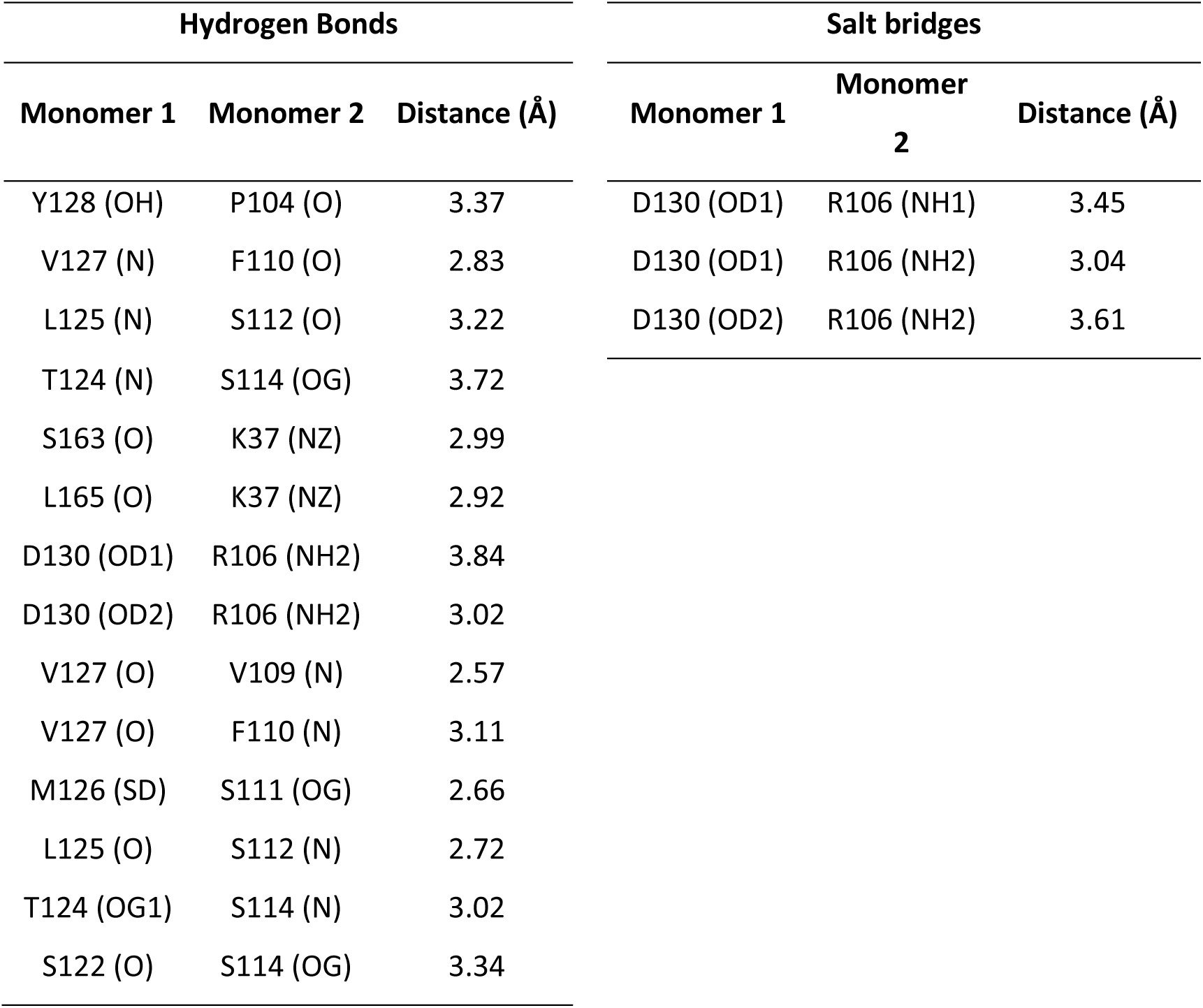
Residues involved in the FAM171A1 trimer interface. Analysis using PDBePISA.

**Supplementary Table 3.** Residues involved in the FAM171A1 trimer-trimer interface. Analysis using PDBePISA.

| FAM171A1 trimer-trimer interfacing residues |  |  |  |
| --- | --- | --- | --- |
| Interfacing residues (buried area %) | Hydrogen Bonds |  |  |
|  | Monomer 1 | Monomer 2 | Distance (Å) |
| Q60 (30) |  |  |  |
| H148 (30) | Y241 (OH) | N239 (OD1) | 3.04 |
| Q150 (20) | Y225 (OH) | K256 (O) | 3.55 |
| R152 (40) | L259 (N) | L259 (O) | 3.82 |
| L187 (40) | L187 (O) | Y225 (OH) | 3.67 |
| D188 (20) | K256 (O) | Y225 (OH) | 3.53 |
| G189 (50) |  |  |  |
| G222 (30) |  |  |  |
| P223 (70) |  |  |  |
| Y225 (70) |  |  |  |
| N239 (30) |  |  |  |
| Y241 (60) |  |  |  |
| R246 (10) |  |  |  |
| L251 (80) |  |  |  |
| L255 (80) |  |  |  |
| K256 (60) |  |  |  |
| S257 (80) |  |  |  |
| G258 (90) |  |  |  |
| L259 (90) |  |  |  |
| G260 (100) |  |  |  |
| L261 (80) |  |  |  |
| T272 (10) |  |  |  |
| Y273 (40) |  |  |  |
| I274 (90) |  |  |  |
| A275 (60) |  |  |  |
| P276 (60) |  |  |  |
| Q277 (10) |  |  |  |

**Supplementary Table 4.** Residues involved in the FAM171A2s trimer interface. Analysis using PDBePISA.

| FAM171A2s trimer interfacing residues |  |  |
| --- | --- | --- |
| FROM domain 1<br>TO domain 2<br>(buried area %) | FROM domain 2<br>TO domain 1<br>(buried area %) | Interactions at centre<br>of trimer<br>(buried area %) |
| K44 (30) | P126 (90) | E52 (40) |
| Q46 (70) | E127 (20) |  |
| Y48 (60) | R128 (50) |  |
| V49 (60) | P129 (100) |  |
| S50 (10) | A130 (90) |  |
| G51 (30) | T131 (100) |  |
| E52 (40) | L132 (100) |  |
| L53 (80) | I133 (100) |  |
| W92 (20) | L134 (100) |  |
| F102 (20) | Y135 (80) |  |
| V107 (70) | E136 (30) |  |
| P108 (70) | D137 (10) |  |
| W109 (100) | L138 (30) |  |
| R110 (30) | H140 (10) |  |
| V111 (100) | L143 (30) |  |
| D112 (10) | G144 (50) |  |
| K113 (40) | S145 (70) |  |
| L114 (20) | P146 (60) |  |
| P115 (80) | R158 (30) |  |
| L116 (50) | Y169 (40) |  |
| Y117 (90) | S170 (50) |  |
| A118 (80) | Q171 (10) |  |
| S119 (100) | L172 (80) |  |
| V120 (100) | W173 (40) |  |
| S121 (90) |  |  |
| Y123 (70) |  |  |

| **<u>Hydrogen Bonds</u>** |  |  |
| --- | --- | --- |
| **Monomer 1** | **Monomer 2** | **Distance (Å)** |
| Y123 (OH) | P129 (O) | 2.08 |
| S121 (OG) | A130 (O) | 3.56 |
| S121 (N) | T131 (OG1) | 3.59 |
| S119 (N) | L132 (O) | 2.88 |
| L116 (N) | L134 (O) | 2.69 |
| Y117 (N) | L134 (O) | 3.54 |
| L44 (NZ) | S170 (O) | 3.82 |
| Y117 (OH) | S170 (O) | 3.65 |
| Y117 (OH) | L172 (O) | 3.31 |
| P108 (O) | S145 (OG) | 3.13 |
| Y117 (O) | L134 (N) | 2.91 |
| S119 (O) | L132 (N) | 2.98 |

